# Modulation of Siglec-7 Signaling via *in situ* Created High-affinity *cis*-Ligands

**DOI:** 10.1101/2020.07.15.203125

**Authors:** Senlian Hong, Chenhua Yu, Shi Yujie, Peng Wang, Digantkumar G. Chapla, Emily Rodrigues, Kelly W. Moremen, James C. Paulson, Matthew S. Macauley, Peng Wu

**Affiliations:** Department of Molecular Medicine, The Scripps Research Institute, La Jolla, California, 92037, U. S. A.; School of Medicine, Nankai University, Tianjin 300071, China; Complex Carbohydrate Research Center, University of Georgia, Athens, Georgia, 30602, U. S. A.; Departments of Chemistry, Medical Microbiology and Immunology, University of Alberta, 11227 Saskatchewan Dr. NW, Edmonton, AB T6G 2G2, Alberta, Canada

## Abstract

Sialic acid-binding immunoglobulin-like lectins, also known as Siglecs, have recently been designated as glyco-immune checkpoints. Through their interactions with sialylated glycan epitopes overexpressed on tumor cells, inhibitory Siglecs on innate and adaptive immune cells modulate signaling cascades to restrain anti-tumor immune responses. However, the mechanisms underlying these processes are just starting to be elucidated. We discover that when human natural killer (NK) cells attack tumor cells, glycan remodeling occurs on the target cells at the immunological synapse. This remodeling occurs through both transfer of sialylated glycans from NK cells to target tumor cells and accelerated *de novo* synthesis of sialosides on the tumor cell themselves. The functionalization of NK cells with a high-affinity ligand of Siglec-7 leads to multifaceted consequences in modulating Siglec-7-regulated NK-activation. At high levels, the added Siglec-7 ligand suppresses NK-cytotoxicity through the recruitment of Siglec-7, whereas at low levels the same ligand triggers the release of Siglec-7 from the cell surface into the culture medium, preventing Siglec-7-mediated inhibition of NK cytotoxicity. These results suggest that glycan engineering of NK cells may provide a means to boost NK effector functions.

## Introduction

The development of immune checkpoint inhibitors for blocking the suppressive functions of cytotoxic T-lymphocyte-associated protein 4 (CTLA-4) and programmed cell death protein 1 (PD-1) has offered curative hopes for many cancer patients.^1-3^ Upon checkpoint blockade treatments, remarkable tumor regression, and significant survival benefits for patients with a broad spectrum of advanced cancers have been observed clinically. However, more than 50% of cancer patients fail to respond to such treatments and some initial responders eventually develop resistance to these therapies with relapsed disease.^2^ The mechanisms leading to such resistance are varied, but one possibility is the existence of other immune checkpoints that are orthogonal to the above well-established ones. Recently, the Siglecs (sialic acid-binding immunoglobulin-like lectins) family of sialic acid-binding proteins has been designated as glyco-immune checkpoints.^4-9^ Through their interaction with sialylated glycan epitopes aberrantly expressed on tumor cells, inhibitory Siglecs such as Siglec-7 and -9 that are found on innate immune cells, e.g. natural killer (NK) cells, inhibit immune signaling pathways to restrain immune-system activation.^6,7,10-12^ Likewise, Siglec-15 upregulated on tumor cells interact with yet-to-be-identified T-cell membrane glycoproteins to suppress T cell anti-tumor functions.^5^ Despite these exciting discoveries that correlate Siglec-mediated immune suppression to cancer progression, we have a poor understanding of the prevalence of the Siglec-ligand expression. Moreover, the mechanisms that govern Siglec-ligand-interaction-mediated immune suppression is only partially elucidated.^7,13,14^

When Siglec-7 or -9 expressing NK cells encounter their target tumor cells, binding to *trans* ligands expressed by tumor cells recruits these Siglecs to the immune synapse, where their immunoreceptor tyrosine-based inhibition motifs (ITIMs) are phosphorylated by Src kinase, thereby creating a binding site for the tyrosine phosphatases SHP-1 and SHP-2.^8,10,15^ Binding of SHP-1/2 leads to dephosphorylation of signaling components downstream of activation receptors to suppress NK cell activation and effector function. Therefore, it seems reasonable to postulate that tumor cells express higher levels of Siglec ligands to counteract NK-induced killing. However, when surveying tumor specimens and matched normal tissues, we observed that high Siglec-7 ligand (Siglec-7L) expression was found in both normal and malignant tissues. Here, we report that when live cancer cells encounter NK cells, a significant upregulation of Siglec-7 ligands is detected within one hour, suggesting that this inhibitory pathway may serve as a negative feedback mechanism that follows NK cell infiltration. Using sialyltransferase-mediated chemoenzymatic glycan editing, a high-affinity Siglec-7 ligand could be newly created on NK cells directly. Introducing such a ligand, in *cis*, induces multifaceted effects on Siglec-7 signaling and NK effector function.

### Siglec-7 ligands are not specifically found on cancer cells and malignant tissues

To understand how the interactions between Siglec-7 and its sialic acid ligands on cancer cells are involved in modulating NK cell-mediated tumor cell killing, we first assessed the expression of Siglec-7 ligands on a panel of cancer cell lines and primary human immune cells using a recombinant Siglec-7 Fc chimera by flow cytometry. Among the cancer cell lines that we screened, Raji B lymphoma and MDA-MB-435 melanoma cells were found to possess the highest level of Siglec-7 ligands (Figure 1a), whereas prostate cancer cell line LNCaP and several ovarian and colorectal cancer cell lines only showed basal levels of Siglec-7 ligands. Although only basal levels of Siglec-7 ligands were found on vascular endothelial cell line HUVEC, freshly isolated NK cells, peripheral blood mononuclear cells (PBMCs), and activated T cells expressed abundant levels of Siglec-7 ligands. To assess *in situ* expression of Siglec-7 ligands, we analyzed paraffin-embedded human tumor and adjacent tissue specimens from a number of malignancies. Significant Siglec-7 Fc staining was detected in almost all specimens analyzed (Figure 1b-d, Supplementary figure s1, and s2); both primary tumor and the matched adjacent tissue specimens exhibited different levels of Siglec-7 Fc staining, which was abolished upon neuraminidase treatment (Figure 1b and 1c). In particular, we examined 177 specimens from lung cancer patients and 20 specimens from normal lung tissues (Figure 1d, 1e, and Supplementary figure s2), and found that approximately 20–40% of lung cancer tissues and 85% of normal lung tissues expressed high levels of Siglec-7 ligands. These observations demonstrate that Siglec-7 ligands are not specifically found on cancer tissues, suggesting that their presence may not be a reliable marker for cancer cells.

**Figure 1.**
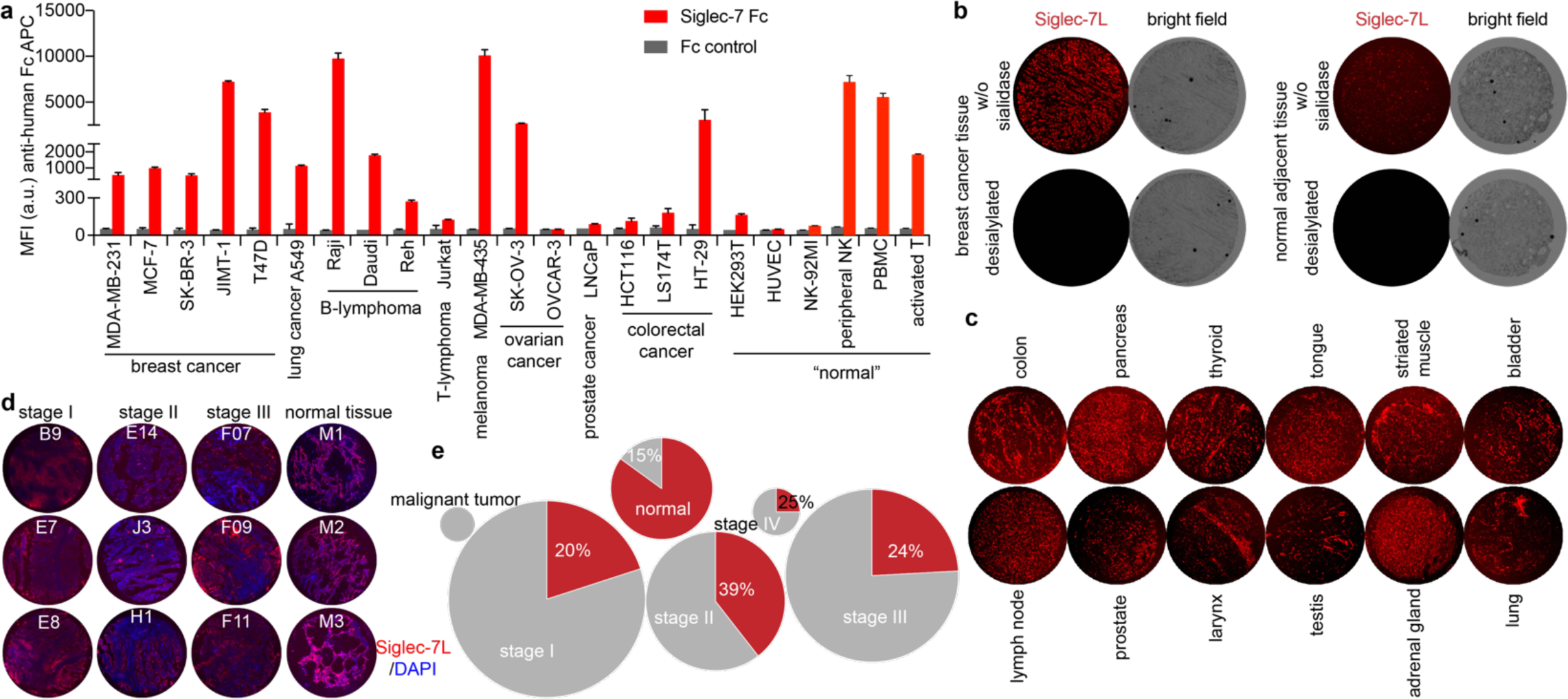
Profile of Siglec-7 ligand (Sigle-7L) expression on the cell surface of live cells and human tissue specimens. (a) Cell lines and primary immune cells from healthy human donors were stained with recombinant Siglec-7 Fc pre-complexed with an anti-human Fc-APC antibody, and anti-human Fc-APC only was used to assess the ligand-binding specificity (Fc control). Error bar represents the standard deviation of three biological samples. (b) Probing the Siglec-7L expression in human breast cancer tissues and the normal adjacent tissues that were pretreated with neuraminidase to remove sialic acids or not. (c) Probing the Siglec-7L expression in different human malignant tissues. (d and e) Probing the Siglec-7L expression in lung cancer tissues and normal tissues from different donors (total of 197 samples). The lung malignant tumor samples (n=2), stage I lung malignant (n=75), stage II lung malignant (n=38), stage III lung malignant (n=58), stage IV lung malignant (n=4), and normal lung tissue (n=20) were used for Siglec-7L screening, and sample size in each group was plotted as circle area (e). Three of the presentative Siglec-7L-high samples were presented (d) Low (gray) and high (red) Siglec-7L levels were classified based on Siglec-7 Fc staining mean fluorescence intensity that determined via ImageJ with a cutoff of 27.00, and the relative ratio of high Siglec-7L specimen was listed out as % percentages..

### Desialylation of Siglec-7L-expressing target cancer cells enhances NK-cell effector functions

Due to the high expression of Siglec-7 ligands on Raji cells, we used these cells as the target (T) cells to determine the impact of the interaction between Siglec-7 on NK cells and cancer cell-expressed Siglec-7 ligands on NK cell effector functions. Primary NK cells^16,17^ isolated from peripheral blood of healthy donors and NK-92MI cells^18,19^ were used as sources of NK effector (E) cells. NK cells isolated from healthy donors were expanded in culture with recombinant human interleukin 2 (IL-2) and IL-15 for 10 days to generate a large number of cells for in *vitro* functional evaluation (these NK cells were defined as peripheral NK cells). The freshly isolated peripheral NK cells contained approximately 40%∼60% of Siglec-7 positive cells, and the frequency of this subset decreased slightly after 10-day *in vitro* culture (Supplementary figure s3). Consistent with previous reports,^20,21^ culture with IL-2 and IL-15 significantly improved the cytotoxicity of peripheral NK cells towards Raji cells (Supplementary figure s3c). To assess the NK-induced target cell killing, we incubated target Raji cells with peripheral NK cells for 4h and determined the target cell killing using a lactate dehydrogenase release assay. As shown in Figure 2a, NK-induced target-cell-killing was enhanced along with increasing the effector cell-to-target cell ratio (E/T ratio). Pre-desialylation of Raji cells by neuraminidase significantly enhanced their susceptibility to the NK-induced killing. Similarly, enhanced Raji cell killing was observed when the Siglec-7 functional blocking antibody s7.7^8^ was used.

**Figure 2.**
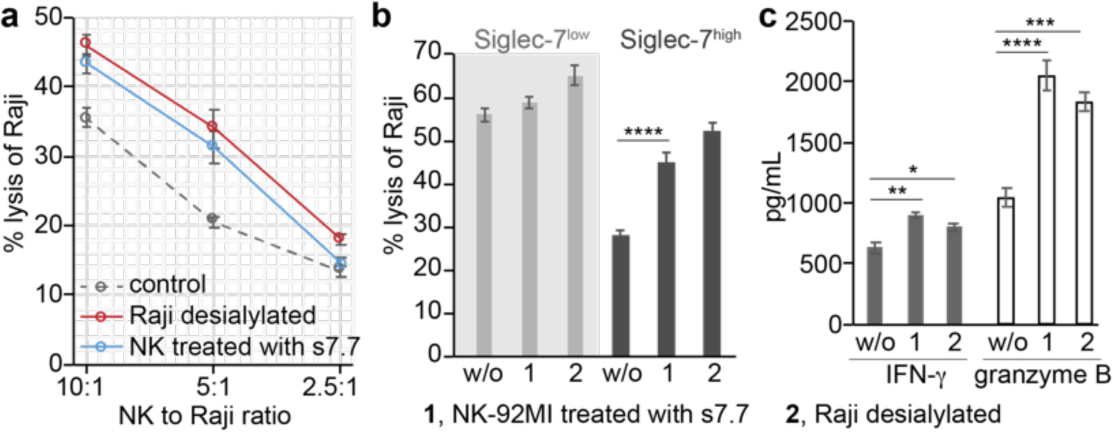
The cytotoxicity of NK cells was impaired by ‘Sialic acid-Siglec-7’ interactions. (a and b) LDH release assay for quantifying the cytotoxicity of the peripheral NK cells (a) and NK-92MI (b) against Raji cells. The pre-desialylation of Raji cells was conducted by neuraminidase treatment. The anti-Siglec-7 antibody (clone s7.7) blocking was performed before the co-incubation of NK with Raji cells. (c) ELISA quantification of IFN-γ and granzyme B produced by NK-92MI-S7^high^ cells after incubation with Raji cells for 1h at an E/T ratio of 5:1. The bars represent the standard error of three biological replicates. The significance was analyzed with the two-sided Student’s t-test. Note, *, p<0.05; **, p<0.01; ***, p<0.005; ****, p<0.001.

Next, we assessed target cell killing using NK-92MI cells, a constantly cytotoxic NK cell line currently undergoing clinical trials as an ‘off-the-shelf therapeutic’ for treating both hematological and solid malignancies.^18,19^ NK-92MI cells were originally derived from a CD56^bright^-NK population,^22^ only a small subset (∼10%) of which expressed Siglec-7 (we define this status as NK-92MI-S7^low^). With increased passages in media supplemented with non-heat-inactivated horse serum, Siglec-7 positive subset became the dominant subset. After approximately 120 days in culture, ∼90% NK-92MI became Siglec-7 positive (we define this status as NK-92MI-S7^high^) (Supplementary figure s4). As expected, compared to NK-92MI-S7^low^ cells, NK-92MI-S7^high^ cells induced approximately 55% reduced killing of Raji cells (E/T ratio = 5:1) (Figure 2b). The NK-92MI-S7^high^-associated cytotoxicity could be enhanced by the Siglec-7 blocking antibody (s7.7) that blocks the ability of Siglec-7 to bind its glycan ligands or by pre-desialylation of Raji cells. Increased IFN-γ and granzyme B secretions were also observed in response to abrogated Siglec-7 ligand interactions (Figure 2c). The same treatments also increased the cytotoxicity of NK-92MI-S7^low^ cells but to a much lower degree (Figure 2b). Similar results were observed using Daudi cells expressing moderate levels of Siglec-7 ligands as the target cells (Supplementary figure s5). Together, these results were consistent with what was reported by the Démoulins group and the Bertozzi group previously,^6,7^ and strongly indicate that Siglec-7 ligands on cancer cells protect them from NK cytotoxicity via interaction with Siglec-7 on NK cells.

### Interactions between NK and tumor cells lead to an upregulation of Siglec-7 ligands on tumor cells

During the course of monitoring interactions between Raji and NK-92MI or peripheral NK cells, we observed a significant increase of sialylated glycans on Raji cell surface within 2 hours, as revealed by lectin staining with *Sambucus nigra* lectin (SNA, specific for α2-6-sialosides) and *Maackia Amurensis* lectin (MAA, specific for α2-3-sialosides) (Figure 3a-c, and Supplementary figure s6). Interestingly, significant increases (about 4fold) of the SNA staining were also observed on JIMT-1 human breast carcinoma and H1975 human non-small cell lung carcinoma cells after incubation with NK-92MI cells under the same condition (Supplementary figure s7). We considered several possibilities through which Raji cells could elevate their surface sialylated glycans: (1) transfer of sialylated glycoconjugates from NK cells, (2) an increase in *de novo* sialic acid synthesis, and/or (3) reduced endocytosis on Raji cell. To examine the first possibility and determine if sialosides on Raji cells were transferred from NK cells, NK cells were cultured in the presence of peracetylated *N*-(4-pentynoyl)mannosamine (Ac_4_ManNAl)^23^ to metabolically incorporate an alkyne-containing Neu5Ac (SiaNAl) onto cell surface glycoconjugates. After 48 hours, NK cells with newly synthesized sialosides labeled by SiaNAl were incubated with Raji cells for 1.5 hour and the cell mixture was reacted with biotin-azide via the ligand (BTTPS)-assisted copper-catalyzed azide-alkyne [3+2] cycloaddition reaction (CuAAC)^24^, followed by flow cytometry analysis. A robust biotinylation signal was detected on NK cells that were cultured with Ac_4_ManNAl. Upon incubating with Raji cells, a 40% decrease in the NK-associated biotin signal was observed, along with the appearance of the biotin signal on Raji cells, suggesting that alkyne-tagged sialic acids were transferred from NK to Raji cells (Figure 3d and 3e).

**Figure 3.**
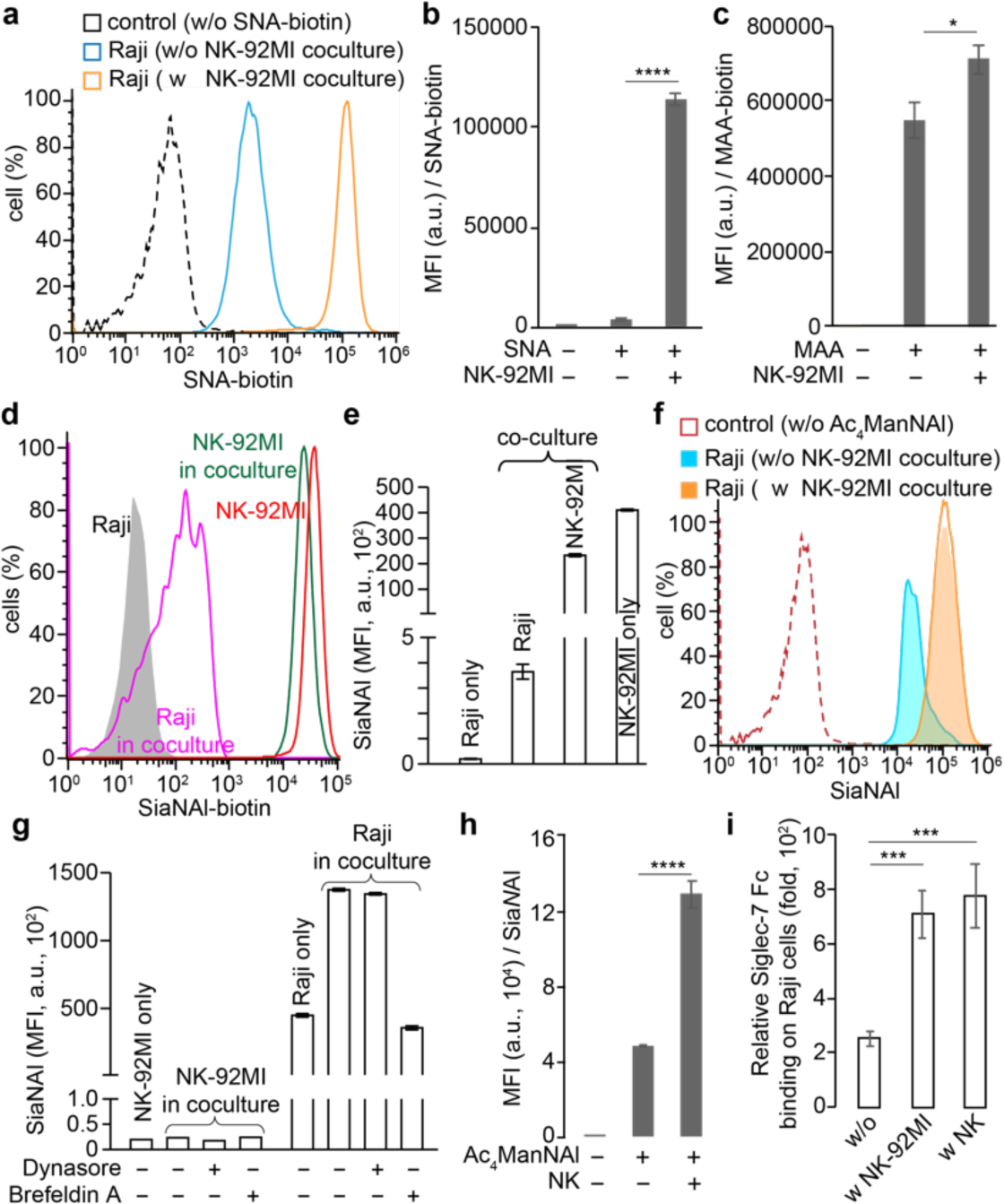
Interactions between NK and Raji cells leads to an upregulation of Siglec-7 ligands on Raji cells. (a) SNA lectin staining of α2-6-sialic acids on Raji cells. (b and c) Mean fluorescence intensity (MFI) of SNA (b) and MAA (c) lectin staining of Raji cells. (d and e) transferring of SiaNAl-labeled sialosides from NK-92MI cells to Raji cells upon the co-culture of both cells. (f and g) Ac_4_ManNAl-based metabolic glycan labeling of Raji cells in the coculture with NK-92MI-S7 cells, in the presence of dynasore, brefeldin A, or not. (h) Coculture with peripheral NK cells also significantly induced Raji cells to express higher sialylation levels, as revealed by Ac_4_ManNAl-labeling. (i) NK-induced up-regulation of sialylation on Raji cells further significantly increased the binding of Siglec-7 Fc. The relative Siglec-7 binding was determined via division to the corresponding Fc controls stained with anti-human Fc-APC. In Figure b, c, e, and g-I, the bars represent standard error of the MFI of three biological repeats of samples. The significance was analyzed with the two-sided Student’s t-test. Note, *, p<0.05; ***, p<0.005; ****, p<0.001.

We next examined the second and third possibilities, whether there was increased *de novo* sialic acid synthesis and/or reduced endocytosis on Raji cells upon NK cell encountering, respectively. Raji cells were metabolically labeled with Ac_4_ManNAl for 48 hours and subjected to the CuAAC-mediated biotin conjugation before and after incubation with NK-92MI cells. In comparison to Raji cells without prior NK incubation, Raji cells that were incubated with NK cells exhibited a 2fold higher biotinylation signal (Figure 3f and 3g). It is noteworthy that the endogenously increased sialylation was at least 200-fold higher than that transferred from NK cells (compare MFI of 400 to 90,000, shown in Figure 3e and 3g). Noteworthy, Raji cells cultured in a contactless, trans-well-based co-culture system together with NK-92MI cells had comparable levels of biotinylation signals as their counterparts that were cultured in fresh and conditional NK-culture medium. This observation indicates that direct NK contact is required for the increase of sialylation on the target cancer cells (Supplementary figure s8). Importantly, we observed that the increased SiaNAl incorporation on the cell surface was largely blocked by Brefeldin A^25^, which inhibits protein transport from the endoplasmic reticulum to the Golgi complex, but not by the endocytosis inhibitor Dynasore^26^ (Figure 3g). Similarly, the co-incubation of Raji cells with peripheral NK cells also triggered a quick accumulation of newly synthesized sialylation, which in turn significantly augmented sialylation on the Raji cell surface (Figure 3h). Notably, no obvious transfer of sialic acids from Raji to NK cells was observed. As a consequence of the increased Raji cell-surface sialylation, a notable elevation of Siglec-7 Fc binding was detected (Figure 3i). In addition, the Siglec-7-specific blocking antibody did not prevent the increase of Raji cell-surface sialylation, suggesting that Siglec-7 may not be involved in triggering this process (Supplementary figure s9). Consistent with a recent study^27^, we found that HEK293T cells (an immortalized human embryonic kidney cell line) that express Siglec-7 ligands on their cell surfaces (Figure 1a), unlike malignant cells, were not attacked by NK cells (Supplementary figure s10). Upon NK encountering, only a slight up-regulation of HEK cell-surface sialylation was detected, and the up-regulated sialic acid was primarily transferred from NK cells rather than *de novo* synthesized (Supplementary figure s10e). Together, the above observations suggest that interactions with NK cells lead to sialoside transfer from the NK cell to the encountered cells. However, d*e novo* sialylated glycan biosynthesis is only induced when the killing of the encountered cells is triggered.

Subsequently, we sought to determine if there is any correlation between NK cell infiltration into tumor tissues and the *in situ* expression of Siglec-7 ligands. We analyzed paraffin-embedded human tumor and adjacent tissue specimens from lung malignancies using the human NK marker CD56 and Siglec-7 Fc staining. The NK density was found to have a modest negative correlation with the Siglec-ligand expression in healthy lung tissue samples (R^2^=0.59), whereas a weak positive correlation existed between NK infiltration and the Siglec-ligand expression in malignant lung tissues (R^2^=0.15) (Supplementary figure s11), suggesting that the up-regulation of Siglec-7 ligands may be triggered by NK cell infiltration into tumor tissues.

### Engineering glycocalyx of NK cells via chemoenzymatic glycan editing

Previous studies by Crocker and coworkers revealed that Siglec-7 on the NK cell surface is masked by *cis* ligands, such as α2,8-linked disialic acids displayed by ganglioside GD3, which partially block its interactions with *trans* ligands found on target cells.^28,29^ Unmasking of Siglec-7 induces strong suppression of NK-induced tumor cell killing due to the binding of Siglec-7 with its *trans* ligands. Based on this precedent, we hypothesized that functionalizing NK cells with a high-affinity ligand of Siglec-7 may further boost NK-associated cytotoxicity by preventing Siglec-7-*trans* ligand interactions. Developed by Paulson and co-workers, ^FTMC^Neu5Ac^30^ (**2**), a chemically-functionalized version of Neu5Ac (**1**) whose C9 is substituted with 9-*N*-(1-(5-fluorescein)-1H-1,2,3-triazol-4-yl)methylcarbamate, when α2-6- or α2-3 linked to *N*-acetyllactosamine (type 2 LacNAc, Galβ1-4GlcNAc), serves as a high-affinity ligand of Siglec-7. We hypothesize that ^FTMC^Neu5Ac could be introduced onto NK cells via sialyltransferase-mediated chemoenzymatic glycan editing to directly create high-affinity Siglec-7 ligands in *cis* for modulating Siglec-7 signaling by preventing its interactions with *trans* ligands.

To assess this feasibility, we first synthesized CMP-^FTMC^Neu5Ac (**5**) as the donor substrate. A commonly used CMP-Sia synthetase from *Neisseria meningitides* (NmCSS)^31^ was not capable of directly converting ^FTMC^Neu5Ac to the corresponding CMP-Neu5Ac analog. Alternatively, we prepared C9-*N*-propargyloxycarbonyl Neu5Ac (**3**, C9-^CPg^Neu5Ac)^32^ and converted it to CMP-C9-^CPg^Neu5Ac. The resulting CMP-C9-^CPg^Neu5Ac (**4**) was then reacted with 5-fluorescein azide^30^ to produce CMP-^FTMC^Neu5Ac (**5**) with an overall yield of ∼26% (Figure 4a and Supplementary figure s20).

**Figure 4.**
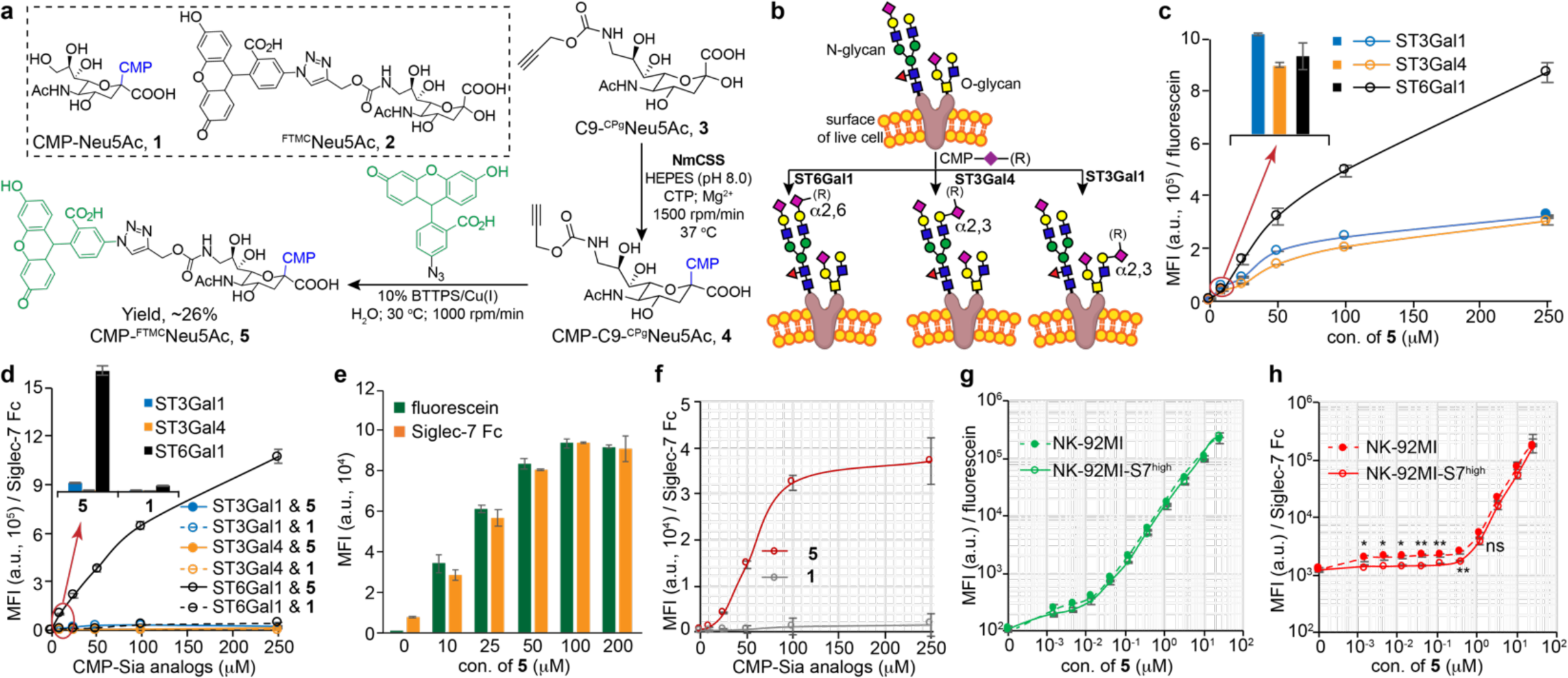
ST-assisted *in situ* creation of Siglec-7 high-affinity ligands on live cells. (a) One-pot synthesis of CMP-^FTMC^Neu5Ac (**5**). (b) Workflow for ST-mediated glycan editing. (c and d) STs-assisted incorporation of ^FTMC^Neu5Ac onto the Lec2 cells was probed with the resultant fluorescein signals (c) and Siglec-7 Fc binding (d). (e) ST6Gal1-assisted incorporation of ^FTMC^Neu5Ac onto the peripheral NK cells was probed with the resultant fluorescein signals and Siglec-7 Fc binding. (f) ST6Gal1-assisted incorporation of ^TMC^Neu5Ac or natural Neu5Ac onto NK-92MI cells was probed by Siglec-7 Fc. (g and h) ST6Gal1-assisted incorporation of ^FTMC^Neu5Ac onto the NK-92MI or NK-92MI-S7^high^ cells was probed with the resultant fluorescein signals (g) and Siglec-7 Fc binding (h). In Figure c-h, the bars represent the standard error of three biological repeats of samples. The significance was analyzed with the two-sided Student’s t-test. Note, ns, not significant; *, p<0.05; **, p<0.01.

Chinese hamster ovary (CHO) Lec2 mutant cells^33^ were used as a model system to test sialyltransferase-mediated chemoenzymatic glycan editing^34^ for incorporating ^FTMC^Neu5Ac onto the cell surface. Lec2 cells possess relatively homogeneous peripheral N-glycans terminated with LacNAc that are capable of being modified by ST6Gal1 and ST3Gal4 to add α2-6- and α2-3-linked sialic acid, respectively (Figure 4b).^35^ Because ^FTMC^Neu5Ac contains fluorescein, cell-associated fluorescein fluorescence can be measured and used to quantify the number of ^FTMC^Neu5Ac installed. After incubating Lec2 cells with ST6Gal1 and ST3Gal4, respectively, in the presence of **5**, a dose-dependent increase of fluorescein signals was detected (Figure 4c). In control experiments, cells were treated with ST3Gal1^35^ to add ^FTMC^Neu5Ac onto O-glycans in α2-3 linkages. When comparable levels of ^FTMC^Neu5Ac were installed onto the cell surface, α2-6-linked ^FTMC^Neu5Ac showed a significantly higher affinity to Siglec-7 than its α2-3-linked counterparts (Figure 4d). For example, whereas similar levels of fluorescein signals were detected upon treating Lec2 cells with each sialyltransferase in the presence of 10 μM CMP-^FTMC^Neu5Ac, ST6Gal1-treated cells exhibited a 14-fold and 75-fold stronger Siglec-7 Fc binding than cells that were treated with ST3Gal1 and ST3Gal4, respectively. These observations indicate that Siglec-7 prefers α2-6-linked ^FTMC^Neu5Ac over its 2-3 linked counterparts (Figure 4c and 4d).

Next, we assessed the efficiency of ST6Gal1-mediated glycan editing on peripheral NK and NK-92MI cells. Consistent with what we observed for Lec2 CHO cells, a dose- and time-dependent increase of fluorescein signals was detected upon treating NK cells with ST6Gal1 and **5** (Figure 4e, 4g, and Supplementary figure s12). When 100 μM of **5** was used for cell-surface glycan editing, Siglec-7 Fc binding reached the saturation level (Figure 4e and 4f). When <1μM of **5** was used for NK-92MI cell modification, we observed an interesting phenomenon: although comparable levels of FTMC-fluorescence could be detected on both NK-92MI Siglec-7^low^ and Siglec-7^high^ cells and fluorescence intensity increased along with increasing dose of **5**, surprisingly, staining these two modified cells with Siglec-7 Fc chimera gave distinct results (Figure 4h). The Siglec-7 Fc staining could be detected on Siglec-7^low^ cells when as low as 10 nM of **5** was used, whereas the staining above background signals on Siglec-7^high^ cells was only detectable when the concentration of **5** went beyond 100 nM. This suggests that the newly installed low-concentration ^FTMC^Neu5Ac may be engaged in strong *cis* interactions with Siglec-7 on NK cell surface, preventing its *trans* interactions with Siglec-7 Fc.

### Modulate NK-induced tumor-cell killing via a newly created *cis* high-affinity Siglec-7L

After confirming that ^FTMC^Neu5Ac could be installed directly onto NK cells to form high-affinity Siglec-7 ligands, we explored the possibility of using this approach to modulate Siglec-7 immunoinhibitory signaling. To our surprise, the killing of Raji cells was significantly inhibited when NK-92MI-S7^high^ cells were functionalized with ^FTMC^Neu5Ac when a high concentration of **5** (i.e. > 100 μM) was used as the donor (Supplementary figure s13). Similar inhibition effects were also observed when desialylated target Raji and Daudi cells were used (Figure 5a and 5b). The decreased cytotoxicity of modified-NK-92MI cells toward desialylated Raji and Daudi cells was gradually rescued when the concentration of **5** used for glycan editing dropped below 1 μM. The lower the CMP-^FTMC^Neu5Ac concentration became the higher the NK-induced target cell killing. The NK cytotoxicity was fully rescued when the concentration of **5** dropped below 10 nM. Remarkably, compared to non-treated NK-92MI cells, NK-92MI-S7^high^ cells modified with 10 nM of **5** exhibited significantly enhanced killing of untrimmed Raji and Daudi cells with their natural sialylation intact (Figure 5c). For example, at the effector-to-target ratio of 5:1, the NK-induced target-cell killing was almost doubled upon ^FTMC^Neu5Ac installation. Consistently, the cytotoxicity of peripheral NK cells and the fluorescence-activated cell sorting (FACS)-isolated Siglec-7 positive peripheral NK cells was also significantly improved upon ST6Gal1-mediated glycan editing with 10 nM of **5** (Figure 5e and Supplementary figure s14).

**Figure 5.**
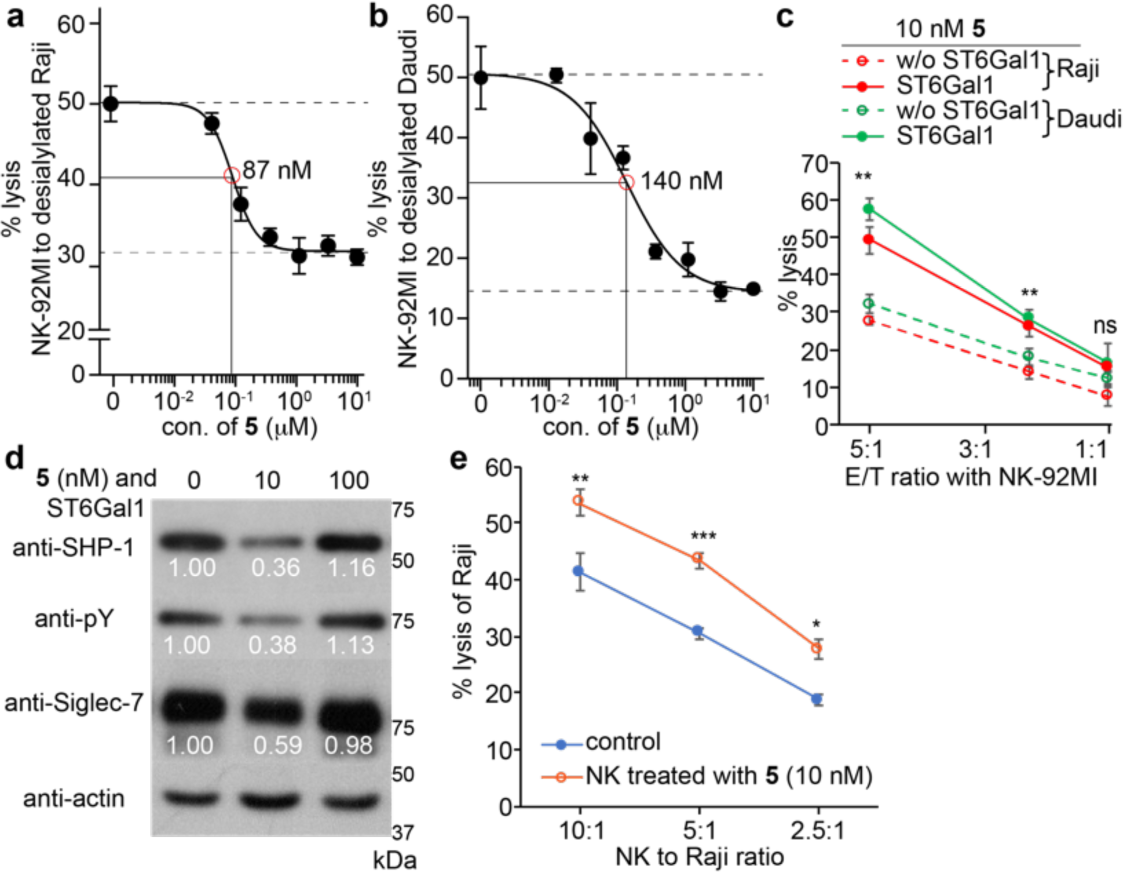
NK cell-surface newly created *cis* high-affinity Siglec-7 ligands modulated NK-cytotoxicity against target cells. (a-c) LDH release assay for quantifying the cytotoxicity of the NK-92MI-S7^high^ cells against Raji (a and c) and Daudi cells (b and c). The NK-92MI-S7^high^ cells with or without ^FTMC^Neu5Ac incorporation were used (d) Western blot analysis of Siglec-7 activation. NK-92MI-S7^high^ cells with or without ^FTMC^Neu5Ac modification were incubated with Raji cells and lysed, followed by anti-Siglec-7 immunoprecipitation. A decrease in SHP-1 recruitment and Siglec-7 phosphorylation was seen in NK cells after treating with 10 nM of **5** but not 100 nM of **5**. The numbers indicate the relative quantification of band intensity to that of actin by ImageJ. (e) The specific lysis of Raji using the sorted Siglec-7 positive peripheral NK cells with or without ^FTMC^Neu5Ac modification. In Figure a-c and e, the bars represent standard error of three biological repeats of samples. The significance was analyzed with the two-sided Student’s t-test. Note, ns, not significant; *, p<0.05; **, p<0.01; ***, p<0.005.

When Siglec-expressing immune cells encounter their target cells, binding to the *trans* ligands that are expressed on target cells recruits Siglecs to the immune synapse, where their ITIMs are phosphorylated by Src kinases that are activated due to activation receptor-ligand ligation, thereby creating a binding site for the tyrosine phosphatases SHP-1 and SHP-2. Binding of SHP-1/2 leads to the de-phosphorylation of signaling components of the activatory receptors to suppress immune cell activation and effector function. Consistent with decreased NK cytotoxicity triggered by the functionalization with high concentrations of **5**, enhanced phosphorylation of Siglec-7 was detected, which was accompanied by increased SHP-1 recruitment (Figure 5d). On the other hand, when NK cells were functionalized with a low concentration of **5** (i.e. 10 nM), significantly reduced phosphorylation of Siglec-7 and SHP-1 recruitment was observed. Simultaneously, a dramatically decreased Siglec-7 expression level was unexpectedly detected in NK cells (Figure 5d).

### Newly created ligands modulate Siglec-7 stability on NK cells

To investigate the mechanism through which high-affinity ligands installed in *cis* on NK cells modulate Siglec-7 inhibitory signaling, we first imaged Siglec-7 distribution on the cell surface of NK-92MI-S7^high^ cells using fluorescence microscopy. As is known for Siglec-2 (CD22) on B cells,^36^ in the absence of *trans*-ligands, Siglec-7 formed abundant clusters on NK-92MI-S7^high^ cells (Figure 6a, 1^st^ panel and Supplementary figure s15-17). After the removal of cell-surface sialic acid by neuraminidase, Siglec-7 clusters broke up (Supplementary figure s13). As expected, when NK-92MI-S7^high^ cells encountered target Raji cells, Siglec-7 was recruited to the E/T interface (Figure 6a, 1^st^ panel).

**Figure 6.**
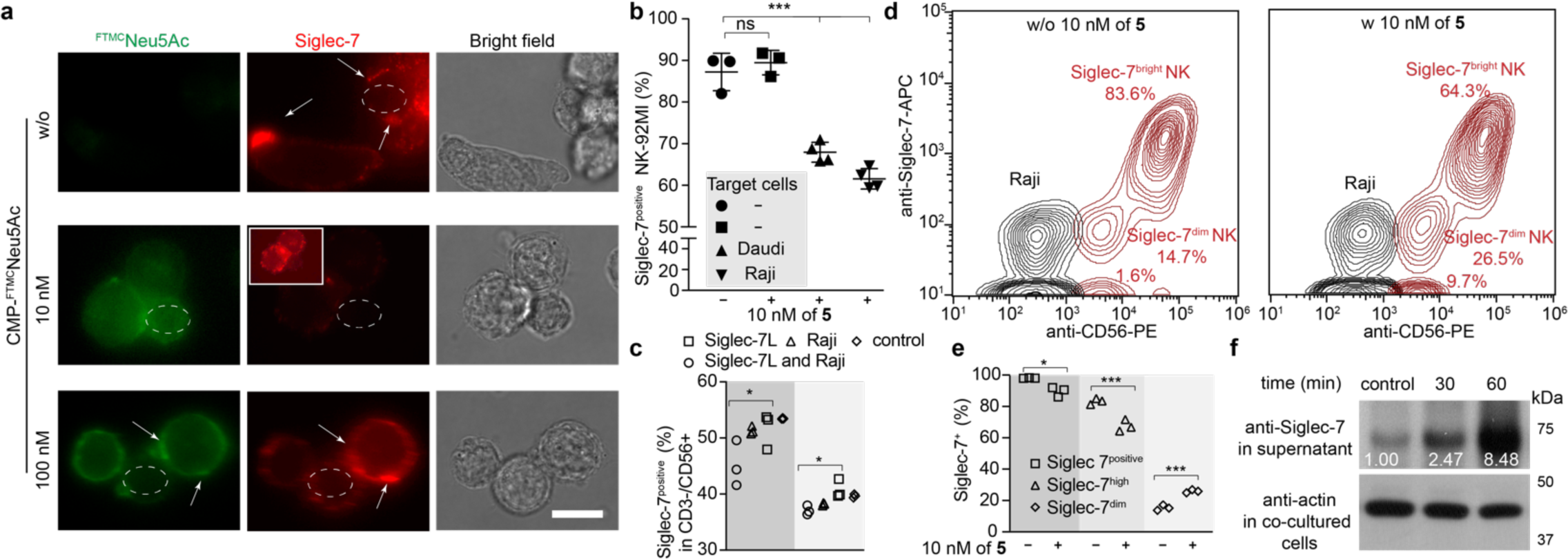
ST6Gal1-assisted incorporation of ^FTMC^Neu5Ac on NK-92MI-S7^high^ or peripheral NK cells modulates Siglec-7 inhibitory signaling. (a) Fluorescence microscopy images of Siglec-7 on NK-92MI-S7^high^ cells with ST6Gal1 or not. The white dash circles indicate the presence of target cells, the white arrows indicate the identified clusters of Siglec-7 and the purple arrows indicate the *in situ* transfer of ligands (green) to target cells from NK cells. White rectangles present the brightened pictures of the same views. Scale bar: 20 μm. (b and c) Siglec-7 positive populations of NK-92MI-S7^high^ cells (b) or peripheral NK cells raised from two different healthy donors (c) treated with ST6Gal1 under 10 nM of **5** were counted via flow cytometry. (d and e) The sorted Siglec-7 positive peripheral NK cells were treated with10 nM of **5** and then were cocultured with Raji cells at an E/T ratio of 5:1. (f) Western blot analysis of Siglec-7 released into the supernatant of the coculture of NK-92MI-S7^high^ cells with Raji cells at an E/T ratio of 5:1. The significance was analyzed with the two-sided Students t-test. Note, ns, not significant; *, p<0.05; ***, p<0.005.

Installation of ^FTMC^Neu5Ac onto the cell surface of NK-92MI-S7^high^ also induced Siglec-7 cluster formation in a dose-dependent manner (Supplementary figure s17). Not only could these clusters be clearly observed on the modified NK cells that were cultured by themselves, but also on the modified NK cells cultured with Raji cells but without direct contact with Raji. Interestingly, upon Raji cell encountering, a dramatic decrease of cell-surface Siglec-7 was detected on NK-92MI-S7^high^ cells that had been modified with 10 nM of **5**, whereas no significant changes of Siglec-7 were observed on NK-92MI-S7^high^ cells that had been modified with >100 nM CMP-^FTMC^Neu5Ac (shown in the 2^nd^ and 3^rd^ panel of Figure 6a)

Consistent with what was observed by imaging, flow cytometry analysis revealed a dramatic decrease (approximately 20∼30%) of the Siglec-7 positive NK cell population following the incubation of target cells with NK-92MI-S7^high^ cells that had been modified with 10 nM of **5** (Figure 6b). Similarly, when encountering with target cells, the Siglec-7 positive subset was decreased among peripheral NK cells (Figure 6c) and the sorted Siglec-7 positive peripheral NK cells (Figure 6d and 6e) that were functionalized with low levels of ^FTMC^Neu5Ac. Surprisingly, no notable increase of Siglec-7 endocytosis was detected as revealed by the weak intracellular Siglec-7 signal. Down-regulation of Siglec-7 expression levels in NK cells (Figure 5d and 6a), coupled with negligible changes in Siglec-7 endocytosis, prompted us to explore the possibility of Siglec-7 release from the plasma membrane. Western blot analysis of Siglec-7 in the culture supernatant revealed that compared to the co-culture supernatant of Raji and un-modified NK-92MI, a dramatically increase of Siglec-7 was detected in the co-culture supernatant of Raji and NK-92MI-modified with 10 nM of **5** (Figure 6f), indicating that Siglec-7 was released from NK plasma membrane upon target cell encountering, a process significantly exacerbated by the installation of super-low levels of ^FTMC^Neu5Ac.

## Conclusion and Discussion

Tumor escape from immune-mediated destruction has been associated with mechanisms that suppress immune system effector functions and trigger immune cell exhaustion.^1-3,37,38^ The Siglec-sialylated glycan interaction is a newly added member of tumor immune escape pathways.^4-8,12-15^ However, the mechanism of this interaction-mediated suppression of anti-tumor immunity is not well understood. Analogous to what has been confirmed for the up-regulated expression of PD-L1 in the tumor microenvironment, which is driven by IFN-γ produced by tumor-infiltrating CD8^+^ T cells,^38^ here, we found that encountering with NK cells triggered up-regulation of Siglec-7 ligands on tumor cells. Therefore, in contrast to initial preconceptions of tumor-intrinsic high expression of Siglec-7 ligands, our data argue that tumor cells upregulate sialylated glycans, which counteract NK-induced killing via the Siglec-sialylated glycan interaction, a mechanism that is intrinsic to the immune system, and likely represents a physiologic negative feedback loop.

Via ST6Gal1-mediated cell-surface glycan editing, we discovered that a high-affinity and specific ligand of Siglec-7 can be created on NK cells to modulate Siglec-7 signaling and NK effector function. With high levels of ligands added onto the NK cell surface, enhanced phosphorylation of Siglec-7 and recruitment of SHP-1 was observed, which in turn suppressed the NK-induced target tumor cell killing. On the other hand, at low levels, the same ligand induced the release of Siglec-7 from the NK cell surface to the culture medium, which completely restored NK cytotoxicity. We confirmed that the ligand-triggered Siglec-7 secretion only took place after the target cell encounter, but not by ligand installation itself. Although chronic HIV infection is known to increase plasma levels of a soluble form of Siglec-7,^40^ to our knowledge, what observed here is the first case that a high-affinity ligand induces the release of Siglec-7 from NK cell surface. It would be interesting to explore whether it is a common phenomenon that functionalizing immune cells with low levels of high-affinity Siglec-ligands could release Siglecs from the cell surface. If so, this strategy may serve as a general approach to down-regulate Siglecs’ inhibitory functions. In this endeavor, we have already demonstrated that high-affinity and specific ligands for other Siglecs, such as Siglec-9^8,30,41^, can be introduced onto live cells by employing similar strategies (Supplementary figure s19 and s21).

## Acknowledgment

This work was supported by the NIH (AI143884 and GM113046 to P.W., P41GM103390, P01GM107012, and R01GM130915 to K.W.M., R01AI050143, P01HL107151, and U19AI136443 to J.C.P.). Prof. M.S.M. would like to thank NSERC, Canada Research Chairs, and the Canadian Glycomics Network for funding. We would like to thank Prof. Mike Boyce for insightful discussions.

## Supplementary Information

**Figure S1.**
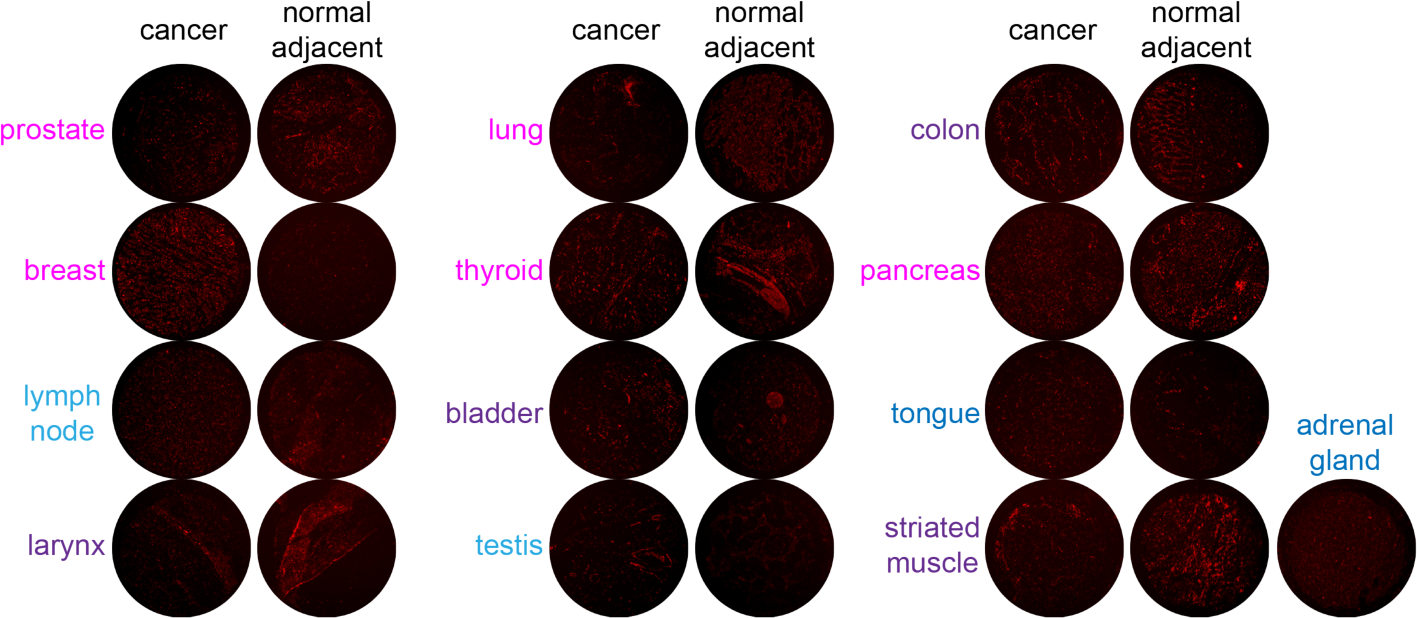
Profile the expression of Siglec-7 ligands on tumor tissues and their related adjacent normal tissues using re-combinant Siglec-7 Fc.

**Figure S2.**
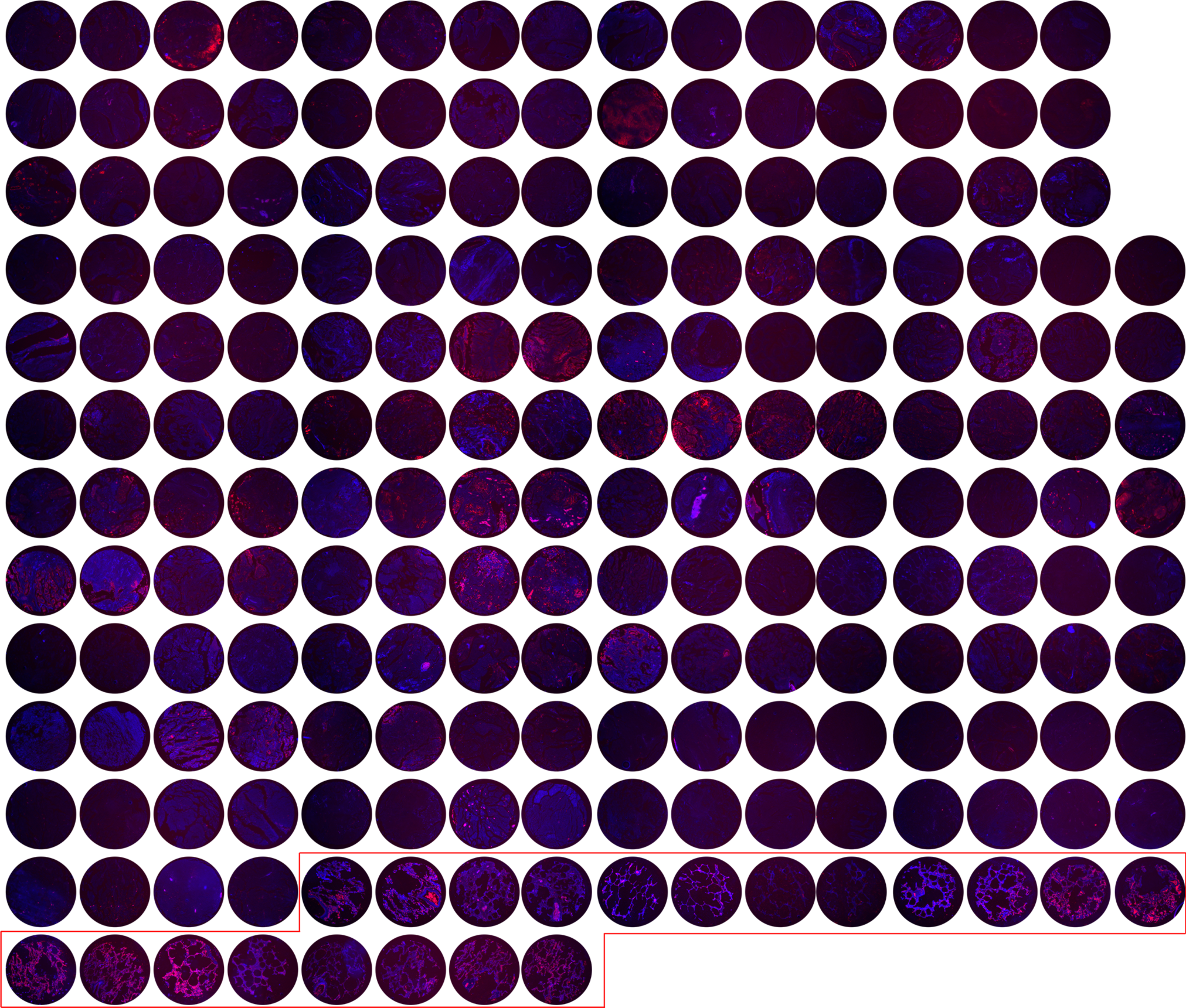
Siglec-7 Fc-based detection of Siglec-7 ligands on lung samples of cancer and healthy tissue (shown in the red box). The cells were stained with DAPI (blue).

**Figure S3.**
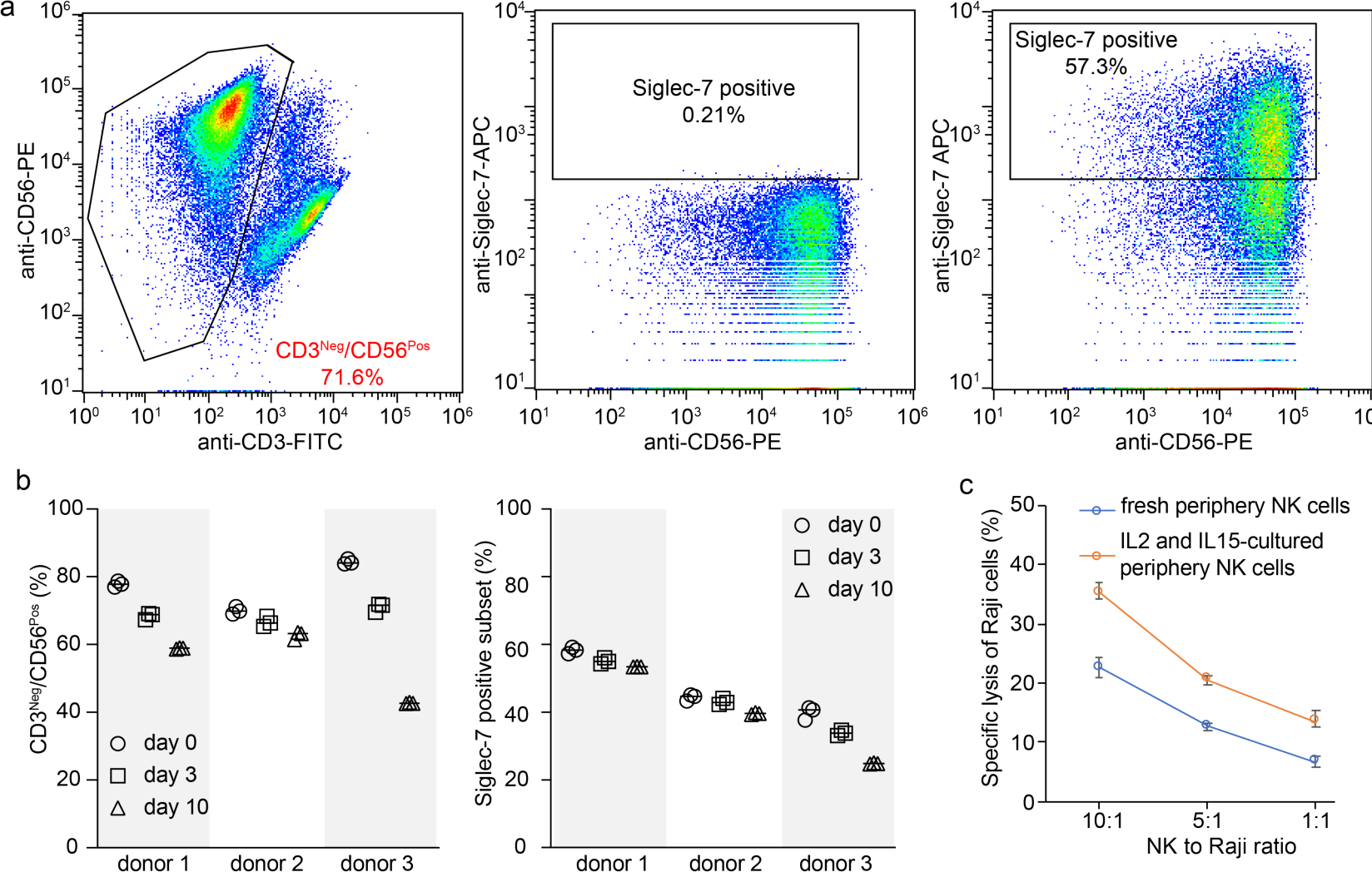
The isolated human periphery NK cells were cultured for 10 days in medium supported with IL-2 and IL-15. (a) The NK cells were gated by anti-CD3 antibody staining (negative) and anti-CD56 antibody staining (positive), and the Siglec-7 expression was probed by anti-Siglec-7-APC (clone 6-434) antibody staining. (b) The CD3^Neg^/CD56^Pos^ and Siglec-7 positive subsets along the culture were monitored by flow cytometry quantification. (c) LDH release assay for quantifying the cytotoxicity of the periphery NK cells against Raji cells. The number of periphery NK cells was determined via CD3^Neg^/CD56^Pos^ population via flow cytometry before incubation with Raji cells at different NK to Raji ratios. Error bars present the standard deviation of three biological repeats.

**Figure S4.**
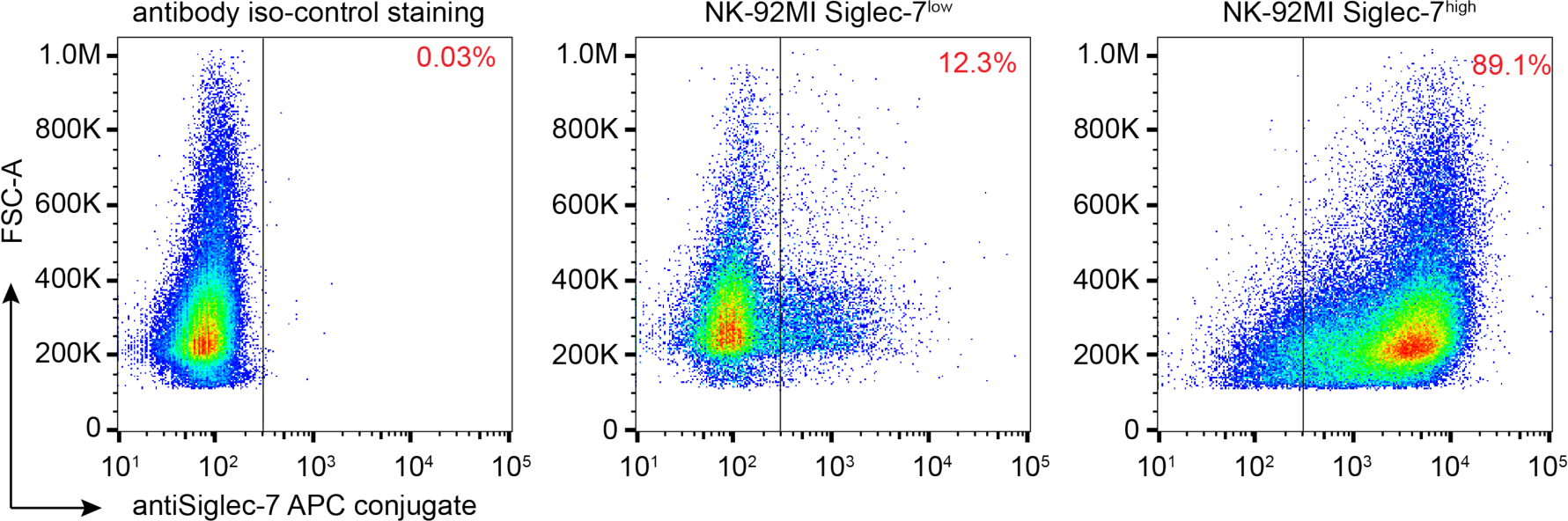
The Siglec-7 expression in NK-92MI cells (NK-92MI-S7^low^) or the sorted NK-92MI-S7^high^ cells were stained with anti-Siglec-7-APC (clone 6-434) antibody and analyzed with flow cytometry. The red number shown in the figure indicates the ratio of the Siglec-7 positive population.

**Figure S5.**
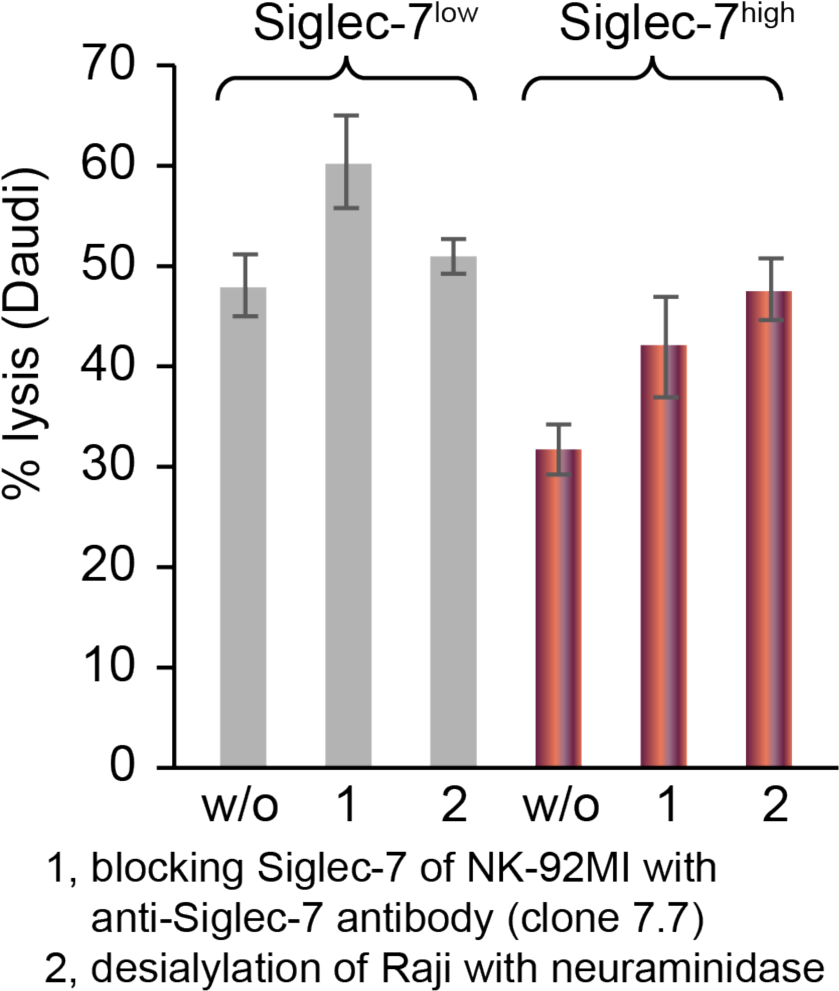
LDH release assay for quantifying the cytotoxicity of NK-92MI-S7^low^ or NK-92MI-S7^high^ cells against Daudi cells, a B-lymphoma cell line with a moderate level of Siglec-7 ligands on the cell surface. The desialylation, ST6Gal1-glycan editing, and anti-Siglec-7 antibody (clone s7.7) blocking were performed before the co-incubation of NK-92MI and Daudi cells at an effector to target ratio of 5:1. Error bars present the standard deviation of three biological repeats.

**Figure S6.**
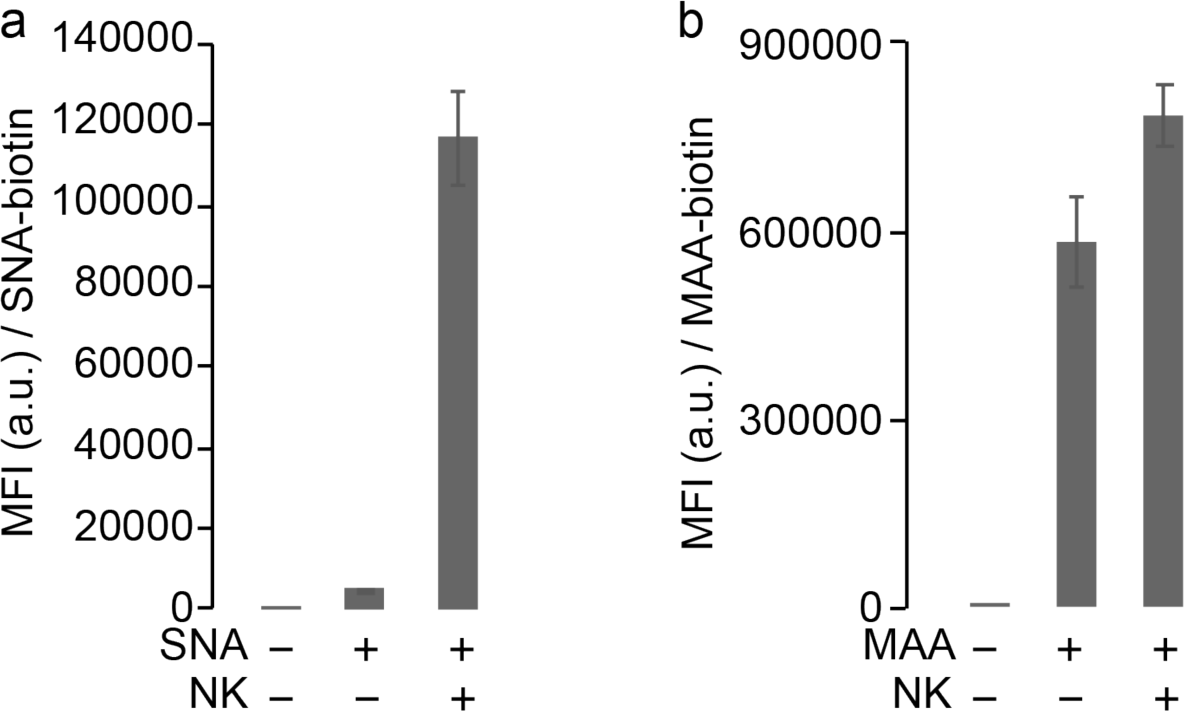
Co-incubation of Raji cells with periphery NK cells significantly up-regulated the level of Raji cell-surface sialylation. The sialosides on Raji cells were probed by SNA (a) and MAA (b) lectin staining. Error bars present the standard deviation of three biological repeats.

**Figure S7.**
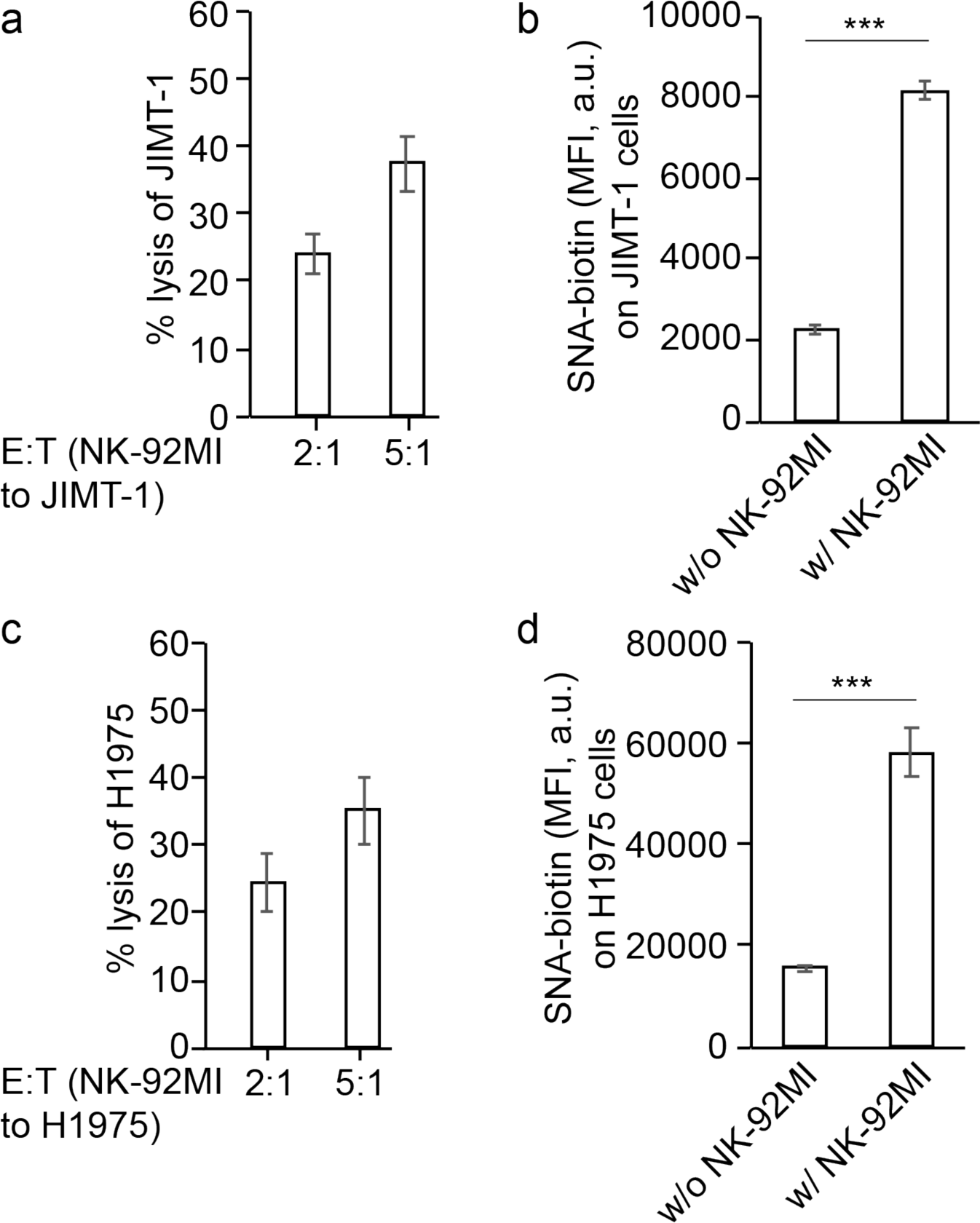
Co-incubation with NK-92MI cells significantly up-regulated the level of cell-surface sialosides on JIMT-1-eGFP-Luc cells (a and b) and H1975-eGFP-Luc cells (c and d). NK-92MI-induced specifc lysis of JIMT-1 (a) and H1975 (b) was quantified via a LDH cytoxicity assay. After Co-incubation of NK-92MI cells with JIMT-1-eGFP-Luc (b) and H1975-eGFP-Luc cells (d) at an E/T ratio of 5:1 for 1.5 h, the sialosides on target cells were stained with SNA-biotin, following by Streptavidin AF647 staining, and flow cytometry analysis. NK-92MI cells in the co-culture system were probed with an anti-CD56 antibody and target cells were marked by the endogenous eGFP fluorescence. Error bars present the standard deviation of three repeats. The significance was analyzed with the two-sided Student’s t-test. Note, ***, p<0.005.

**Figure S8.**
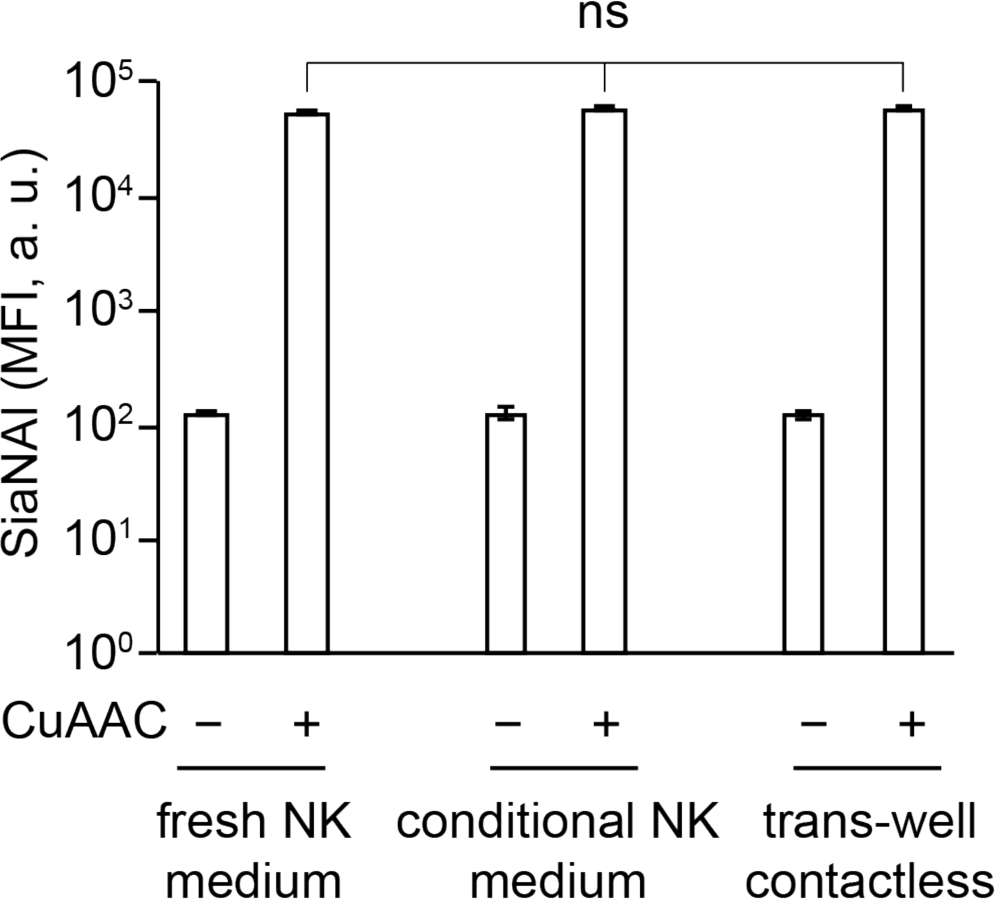
Trans-well based contactless co-culture of NK-92MI cells with Raji cells did not trigger the increase of Raji cell-surface sialosides. Raji cells were cultured with Ac_4_ManNAl for 2 days, and further cultured with fresh NK-92MI complete medium, or conditional medium of NK-92MI, or co-incubation with NK-92MI cells separated via trans-well insertions. After 1.5h, Raji cells were collected for CuAAC-labeling of the metabolically incorporated SiaNAl. Error bars present the standard deviation of three repeats. The significance was analyzed with the two-sided Student’s t-test. Note, ns, not significant.

**Figure S9.**
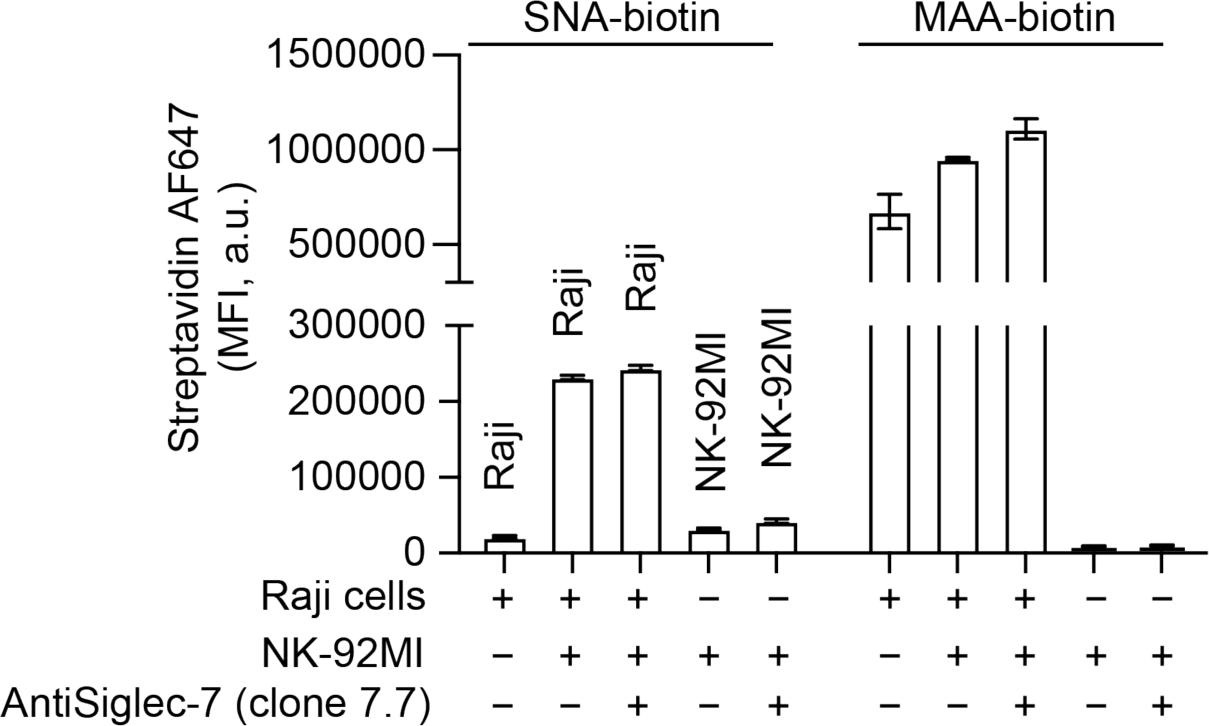
The cell-surface sialylation on Raji cells was increased after co-cultureing with NK-92MI-S7^high^ cells as revealed by lectin staining. NK-92MI and Raji cells cultured alone were stained as controls. NK-92MI-S7^high^ cells in the co-culture system were probed with anti-CD56 antibody and Raji cells were probed by the anti-CD19 antibody. Error bars present the standard deviation of three repeats.

**Figure S10.**
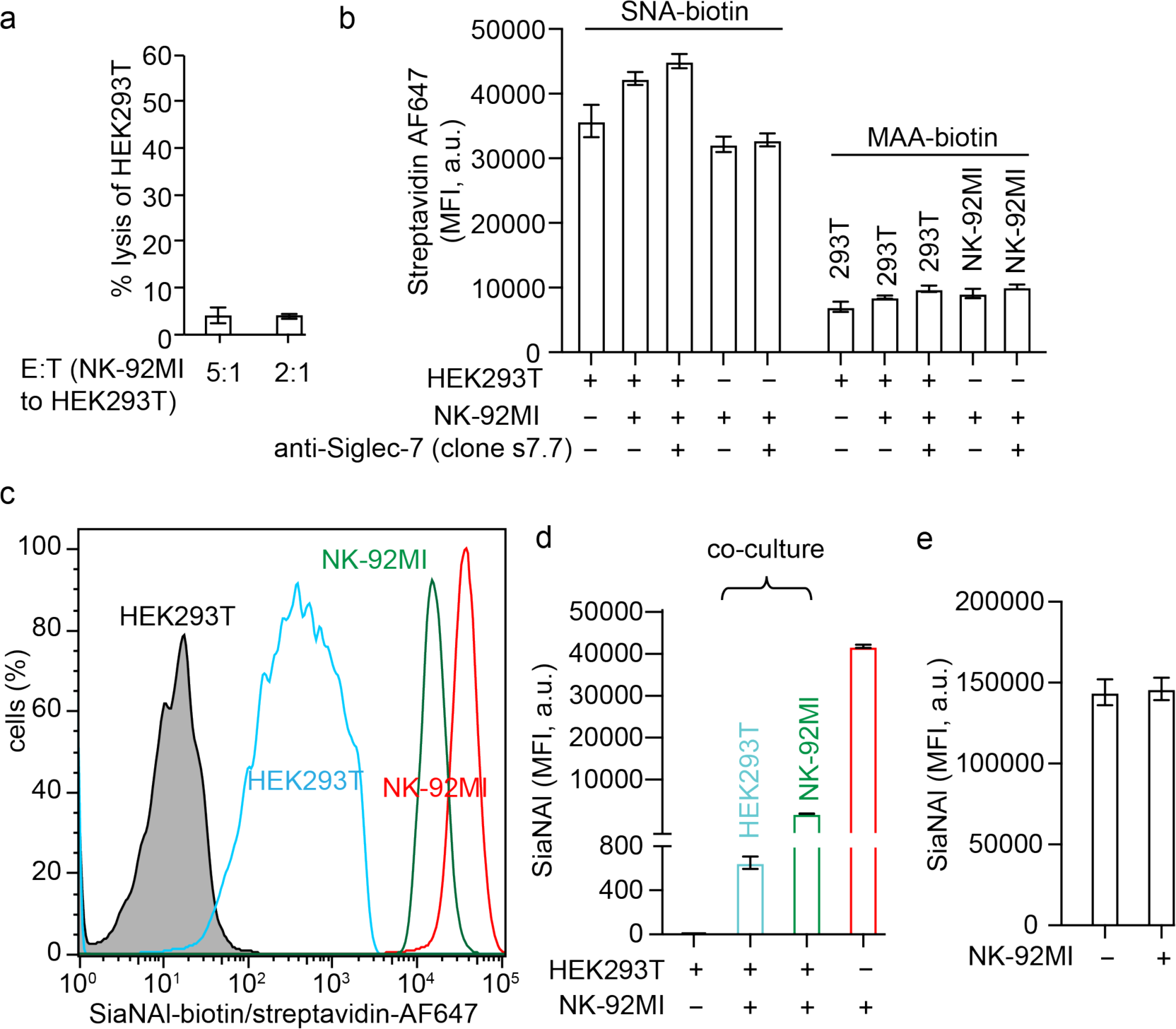
The changes of sialylation on HEK293T cells after encountering NK-92MI-S7^high^ cells. (a) LDH release assay for quantifying NK-92MI-S7^high^ cytotoxicity against HEK293T cells. (b) Probing sialylation on HEK293T cells with SNA and MAA lectins. The NK-92MI-S7^high^ cells (CD56 Positive) in co-culture with HEK293T cells (CD56 Negative) were probed with the anti-CD56 antibody. (c and d) Probing the transfer of NK-92MI-S7^high^ cell-surface sialosides to HEK293T cells via Ac_4_ManNAl-based metabolic glycan labeling. NK-92MI-S7^high^ cell-surface sialosides were metabolically labeled by treating with Ac_4_ManNAl for 2 days. Then, HEK293T cells were coincubated with the treated NK-92MI-S7^high^ cells in fresh medium without Ac_4_ManNAl for 1 h. The resultant SiaNAl-labeled glycoconjugates on HEK293T cells and NK-92MI-S7^high^ cells were further labeled with biotin in DPBS buffer (pH 7.4) containing 2% FBS, 50 μM azide-PEG4-biotin, 300 μM BTTPS and 50 μM Cu^2+^ premixture, and 2.5 mM freshly prepared sodium ascorbate for 5 mins. Finally, cell-surface biotin was probed by 2 μg/mL streptavidin-Alexa Fluor 647 conjugate and quantified by flow cytometry. NK-92MI-S7^high^ and HEK293T cells without co-culture were stained as controls. (e) Probe sialylation biosynthesis in HEK293T cells via Ac_4_ManNAl metabolic labeling. Error bars present the standard deviation of three repeats..

**Figure S11.**
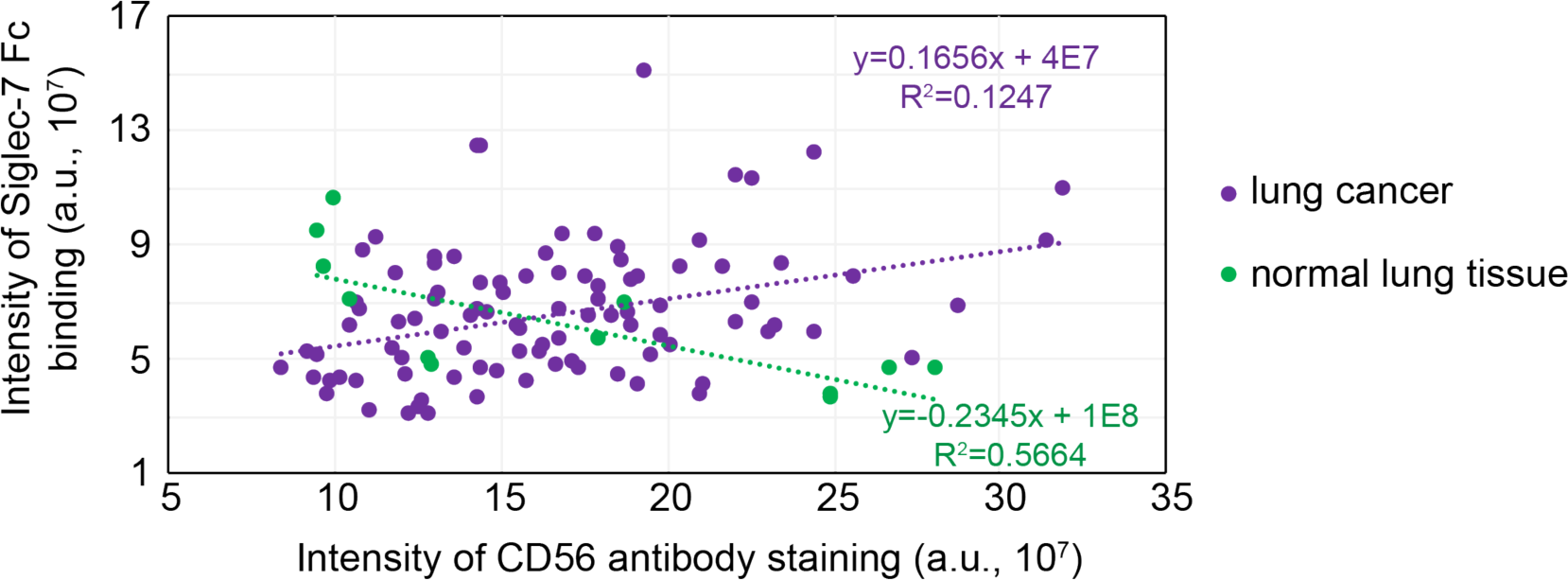
Immunofluorescence analysis of NK marker CD56 and Siglec-7 ligand expression in human lung cancer (n=96) and normal lung (n=12) tissues. Results were obtained using human tissue microarray LC1201a (USBiomax) containing multiple types of lung carcinoma, and probed with anti-CD56 and Siglec-7 Fc, quantification of immunofluorescence via ImageJ and linear correlation analysis in Excel.

**Figure S12.**
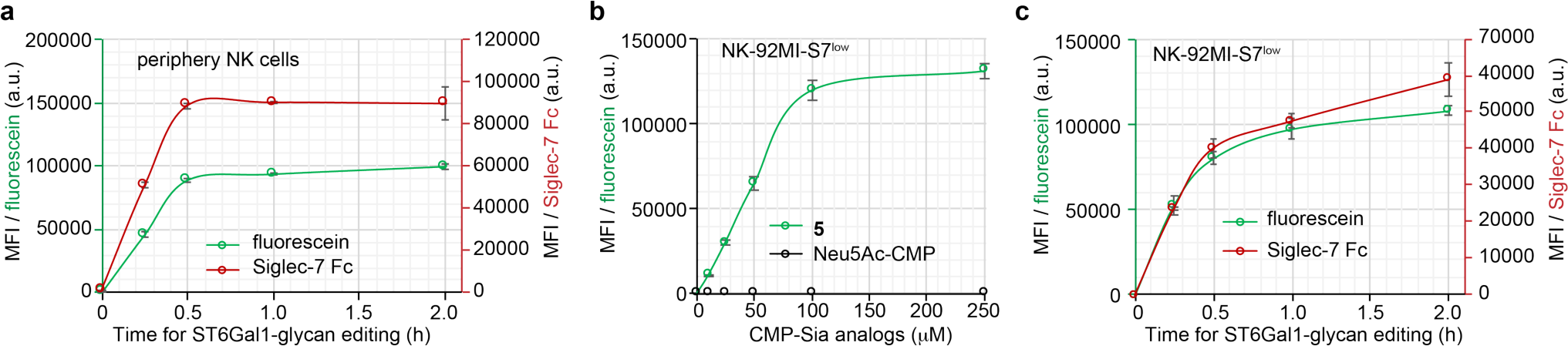
The dose and time-dependent incorporation of CMP-Sia analogs onto periphery NK cells (a) and NK-92MI-S7^low^ cells (b and c) via ST6Gal1-glycan editing. For time-course study (a and c), 200 μM ^FTMC^Neu5Ac-CMP (**5**) and 40 μg/mL ST6Gal1 were used. The error bars present the standard deviation of three repeats.

**Figure S13.**
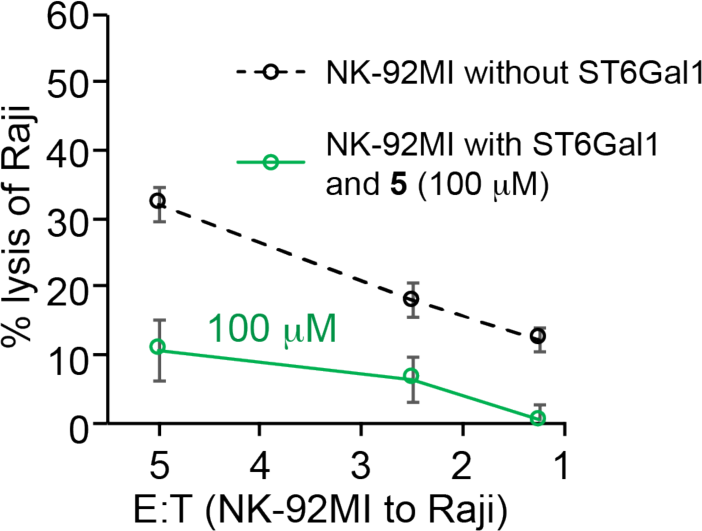
LDH release assay for quantifying cytotoxicity of NK-92MI-S7^high^ cells with ^FTMC^Neu5Ac or not against Raji cells. Before co-incubation with Raji cells for 4 h, the NK-92MI-S7^high^ cells were pre-treated with ST6Gal1and ^FTMC^Neu5Ac-CMP (**5**, 100 μM), or without ST6Gal1 as control. The error bars present the standard deviation of three repeats.

**Figure S14.**
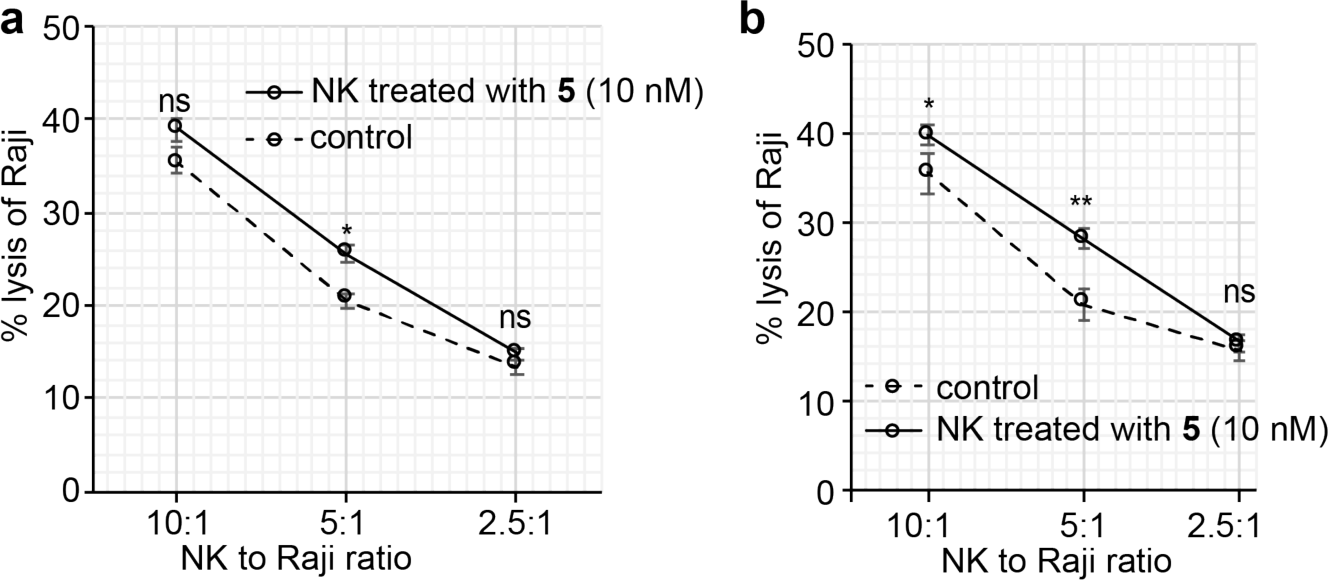
The specific lysis of Raji using the peripheral NK cells with or without ^FTMC^Neu5Ac modification. Raji cells were incubated with peripheral NK cells from two different healthy donors at indicated E/T ratios for 4h. The specific lysis of Raji cells was measured via the LDH release assay. The bars represent the standard error of three biological repeats of samples. The significance was analyzed with the two-sided Student’s t-test. Note, ns, not significant; *, p<0.05; **, p<0.01; ***, p<0.005.

**Figure S15.**
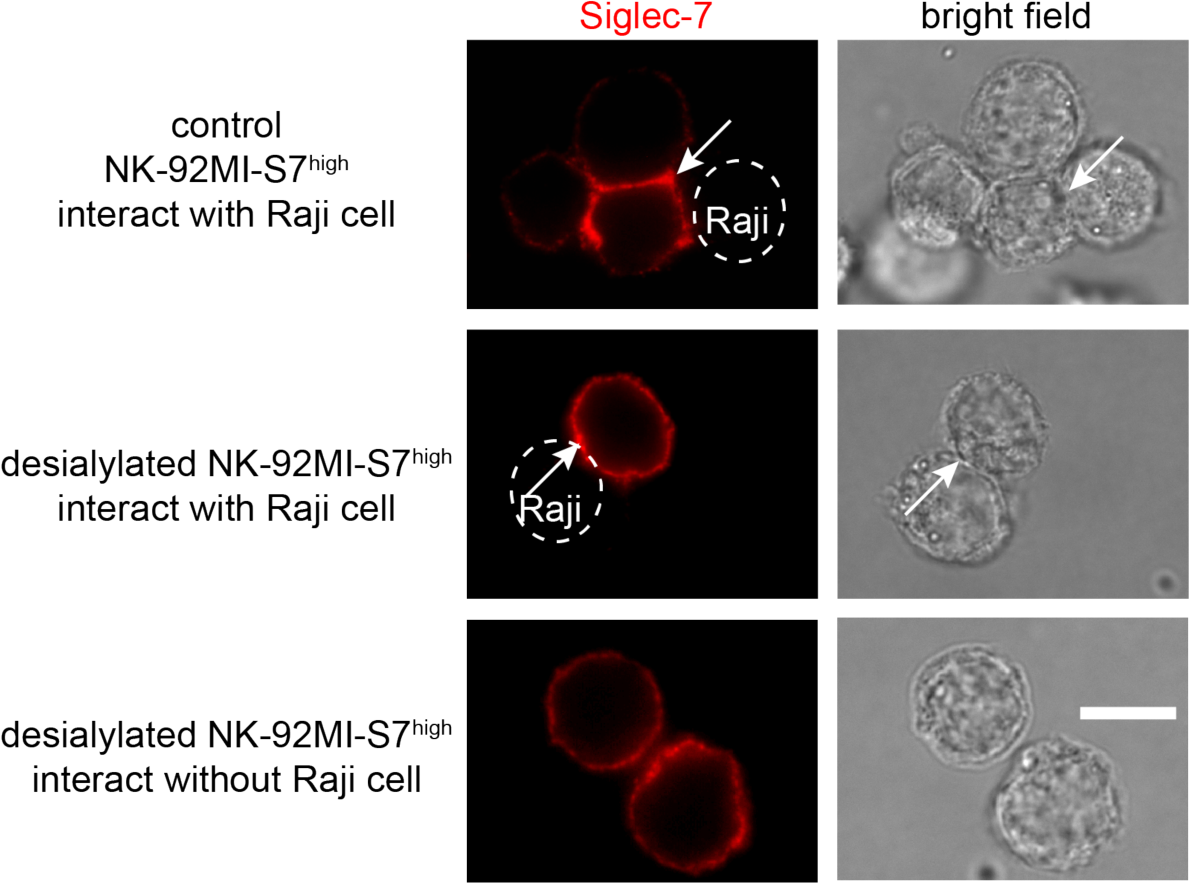
Fluorescent images of NK-92MI-S7^high^ cells with or without ‘self’sialosides after neuraminidase treatment. NK-92MI-S7^high^ cells co-incubated with or without Raji cells were stained with anti-Siglec-7-PE antibody (clone 6-434, in red) and imaged by fluorescent microscopy. NK-92MI-S7^high^ cells desiaylated by neuraminidase before coincubating with target Raji cells at an effector to target ratio of 5:1 for 1 h. The scale bar is 20 μm.

**Figure S16.**
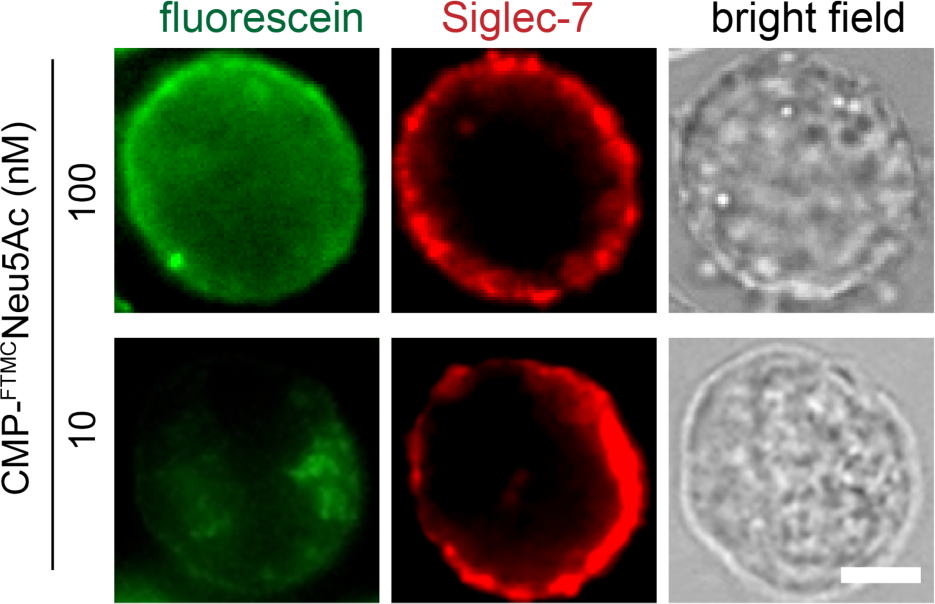
Representative images of NK-92MI-S7^high^ cells modified with ^FTMC^Neu5Ac without interaction with other NK cells or in contact with Raji target cells in the co-culture system. The NK-92MI-S7^high^ cells were treated with ST6Gal1 and CMP-^FTMC^Neu5Ac, and coincubated with Raji cells at an E/T ratio of 5:1. The fluorescein groups on ^FTMC^Neu5Ac were directly imaged (green), and Siglec-7 was probed with anti-Siglec-7-PE. The fluorescein and PE were imaged with a fluorescence microscope. Scale bar: 10 μm.

**Figure S17.**
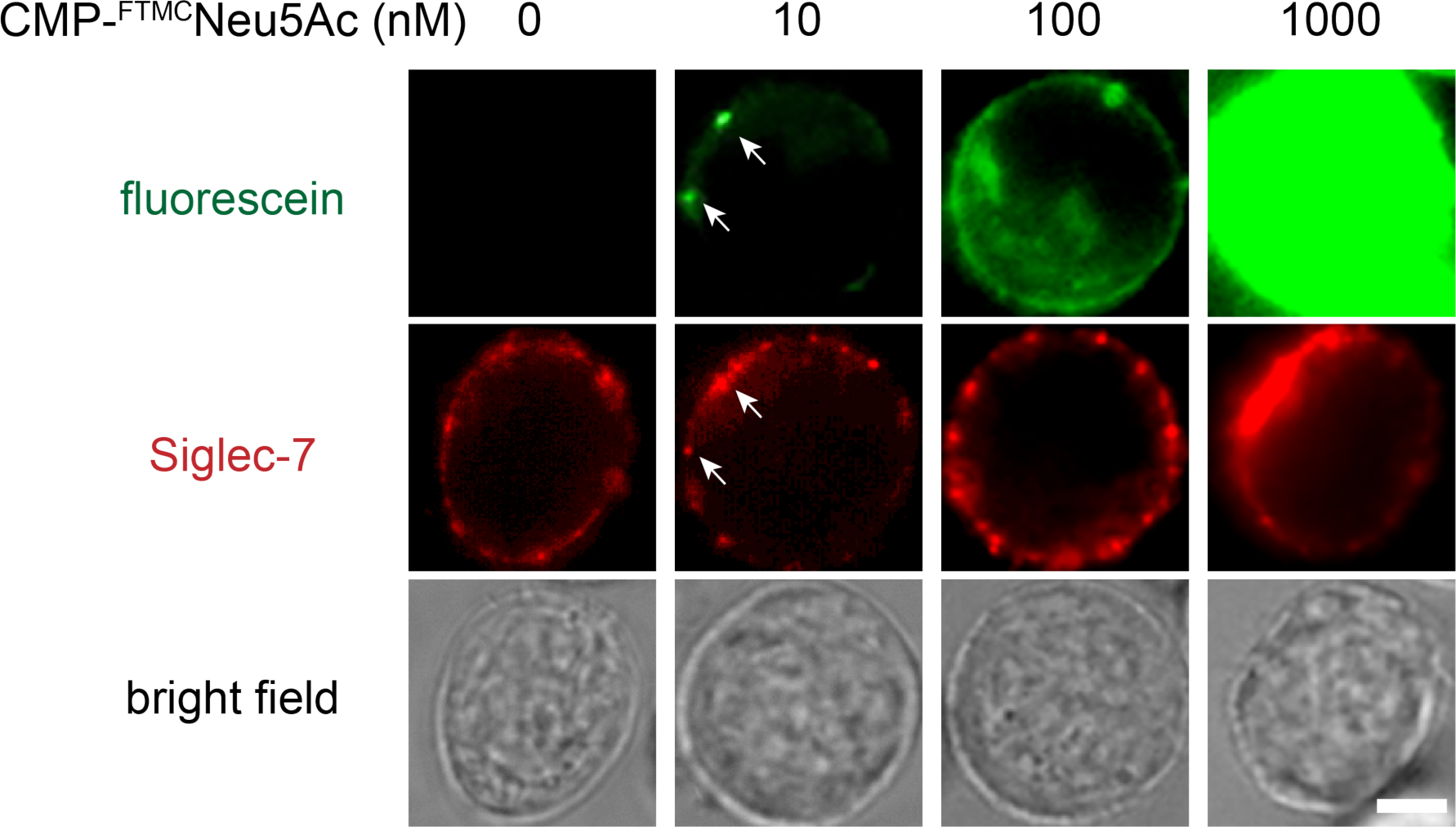
Immunoimaging Siglec-7 on NK-92MI-S7^high^ cells. The NK-92MI-S7^high^ cells were treated with ST6Gal1 and indicated concentrations of CMP-^FTMC^Neu5Ac for 30 min, before the NK cells were washed and stained with anti-Siglec-7-PE antibody. The fluorescein and PE were imaged with a fluorescence microscope. Scale bar: 10 μm.

**Figure S18.**
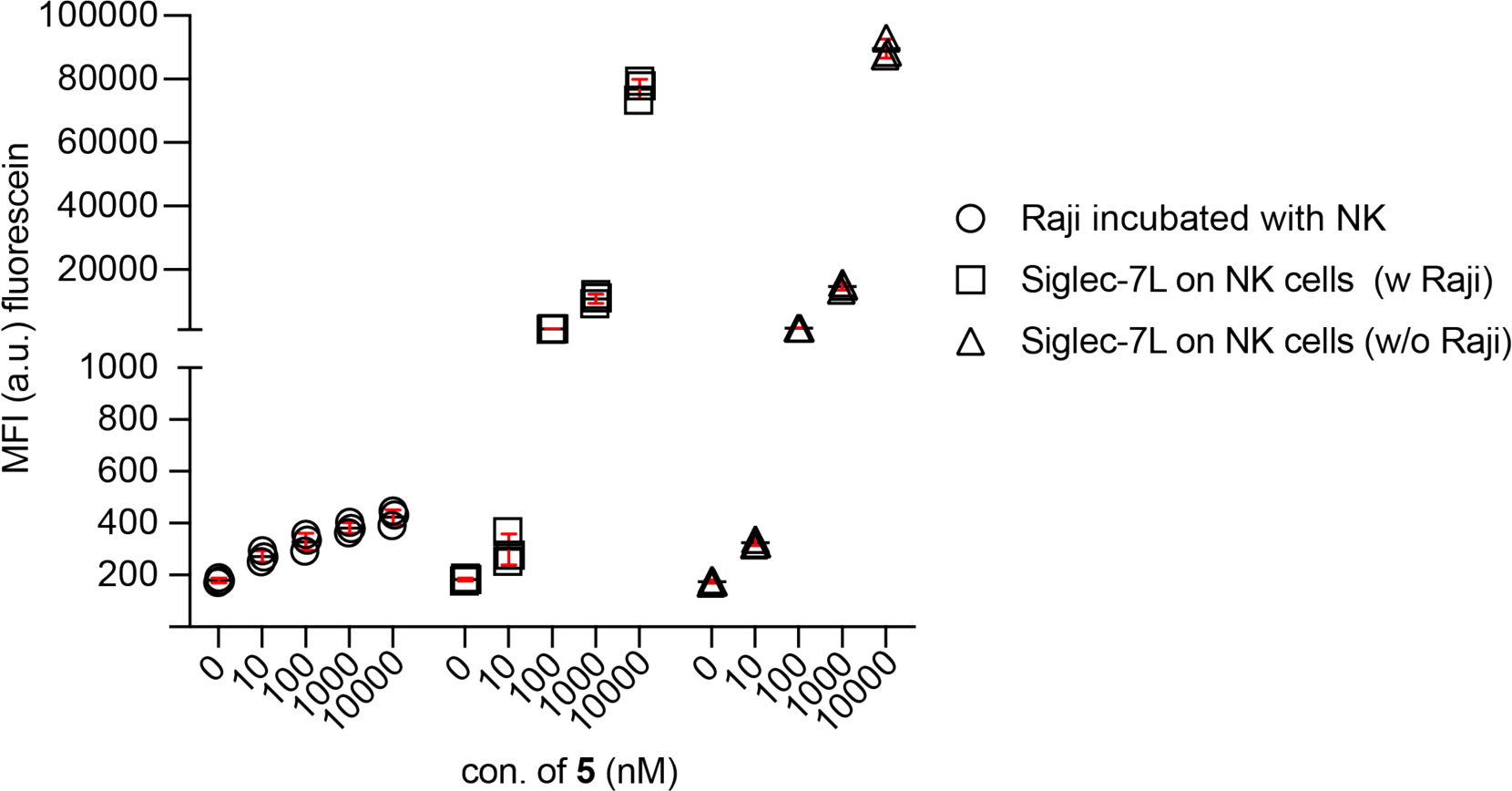
Flow cytometry assisted quantification of high-affinity ligands of Siglec-7 transferred to target Raji cells from ligand-modified NK-92MI cells during coincubation. The NK-92MI cells were treated with ST6Gal1 and indicated concentrations of CMP-^FTMC^Neu5Ac at 37 °C for 30 min. Then the modified or unmodified NK-92MI cells were washed with DPBS before incubation with Raji cells at an E/T ratio of 5:1 for 1 hr. The cell mixture was further stained with anti-CD56-PE and anti-CD19-APC to differentiate Raji and NK-92MI cells. The ligand fluorescein intensity was directly quantified via flow cytometry.

**Figure S19.**
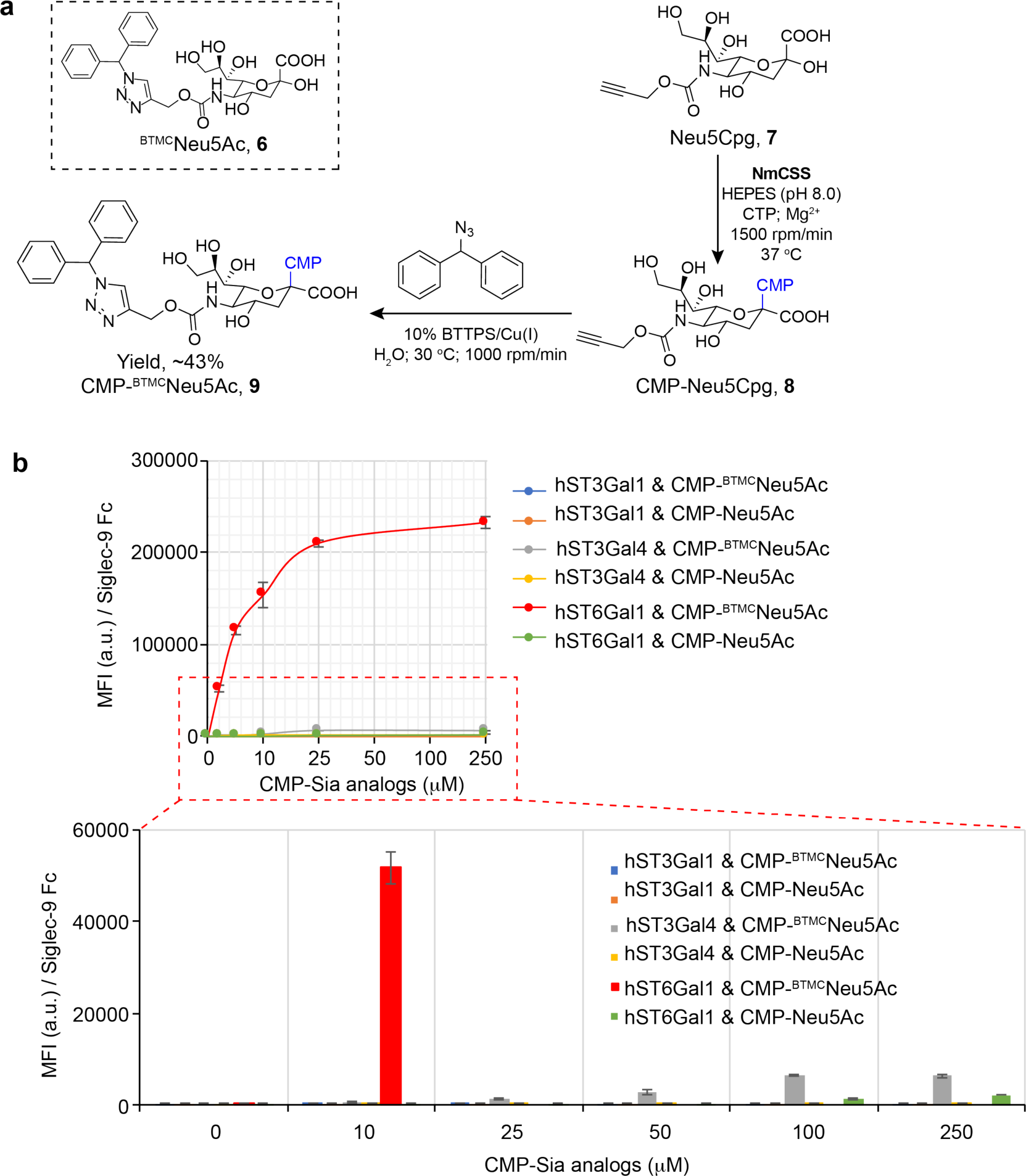
Using in situ creation of Siglec-9 high-affinity ligands to target Siglec-9 signaling. (a) Chemical structure of ^BTMC^Neu5Ac^[23]^ (**6**) and route for one-pot synthesis of CMP-^BTMC^Neu5Ac (**9**). CMP-^BTMC^Neu5Ac (**9**) was prepared from C5-N-propargyloxycarbonyl Neu5Ac (Neu5CPg)^[27]^ (**7**) by conjugating CMP-Neu5CPg (**8**) with diphenylmethyl azide with a yield of ∼43%. (b) STs-assisted incorporation of ^BTMC^Neu5Ac onto Lec2 cells. Lec2 cells treated with STs (ST3Gal1, ST3Gal4, or ST6Gal1) under the presence of CMP-^BTMC^Neu5Ac, CMP-Neu5Ac or not, were stained with Siglec-9 Fc and quantified via flow cytometry. Lower panel figure presents the magnified details of figure upper panel labeled by a red box. Error bars present the standard deviation of three biological repeats.

**Figure S20.**
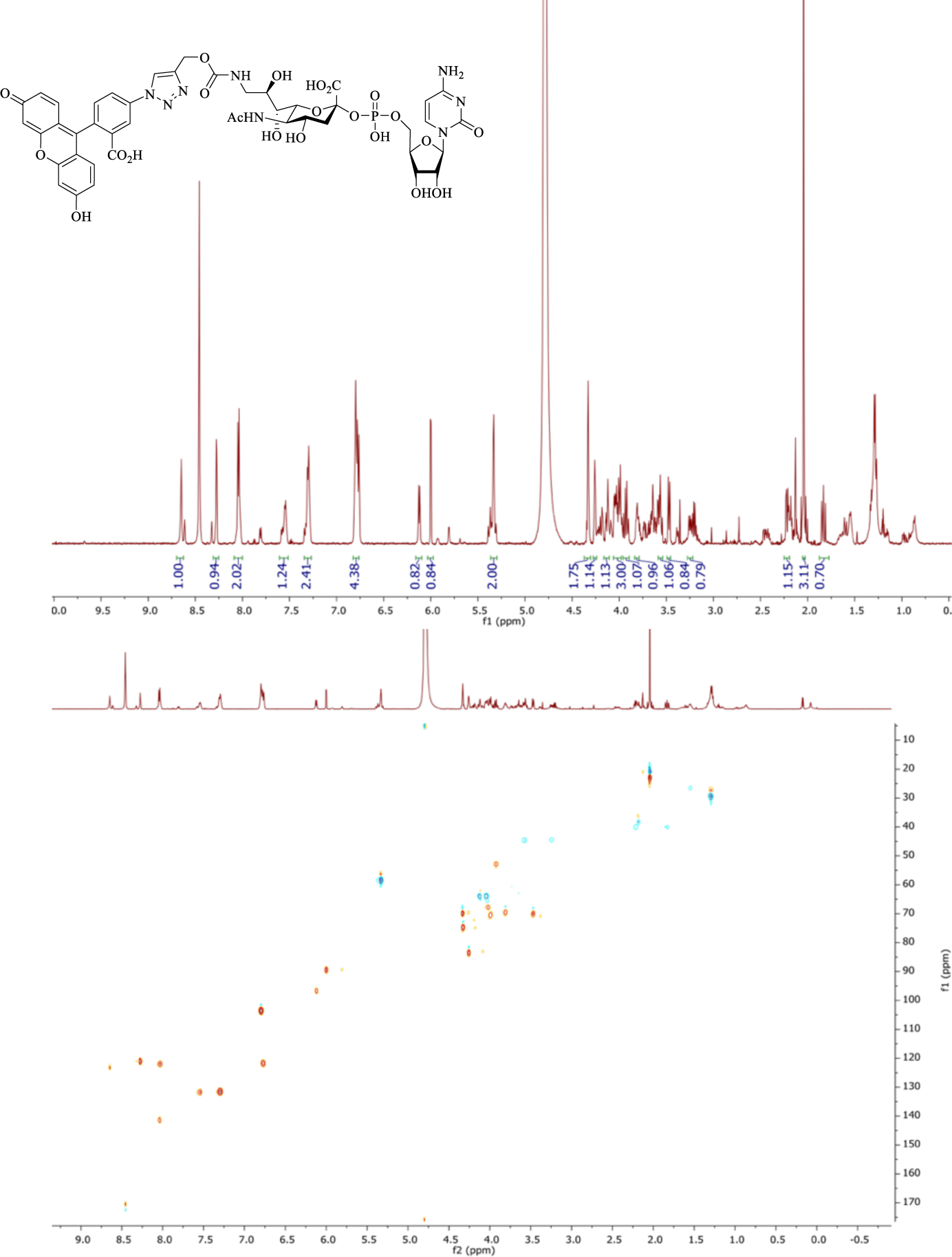
1H and 13C NMR spectra of CMP-^FTMC^Neu5Ac. ^1^HNMR (600 MHz, D_2_O): *δ* = 1.84 (t, 1H, *J* = 12.0 Hz), 2.04 (s, 3H), 2.22 (dd, 1H, *J* = 4.8, 12.6 Hz), 3.24 (dd, 1H, *J* = 7.8, 13.8 Hz), 3.47 (d, 1H, *J* = 8.0 Hz), 3.53-3.58 (m, 1H), 3.78-3.84 (m, 1H), 3.89-3.96 (m, 1H), 3.97-4.06 (m, 3H), 4.10-4.15 (m, 1H), 4.24-4.28 (m, 1H), 4.30-4.37 (m, 2H), 5.29-5.37 (m, 2H), 6.00 (d, 1H, *J* = 3.6 Hz), 6.12 (d, 1H, *J* = 7.8 Hz), 6.74-6.83 (m, 4H), 7.25-7.35 (m, 2H), 7.51-7.61 (m, 1H), 7.99-8.08 (m, 2H), 8.27 (s, 1H), 8.65 (s, 1H). ^13^CNMR (150 MHz, D_2_O): *δ* = 22.82, 40.02, 44.42, 52.6, 58.29, 63.91, 67.75, 69.58, 69.86, 69.95, 70.49, 74.68, 83.47, 89.42, 96.66, 103.46, 121.04, 121.7, 122.08, 123.38, 131.61, 131.78, 141.4 (Observed from HSQC). HRMS: m/z calc. for C_44_H_46_N_8_O_22_P: 1069.2464; found: 1069.2487 [M + H]^+^

**Figure S21.**
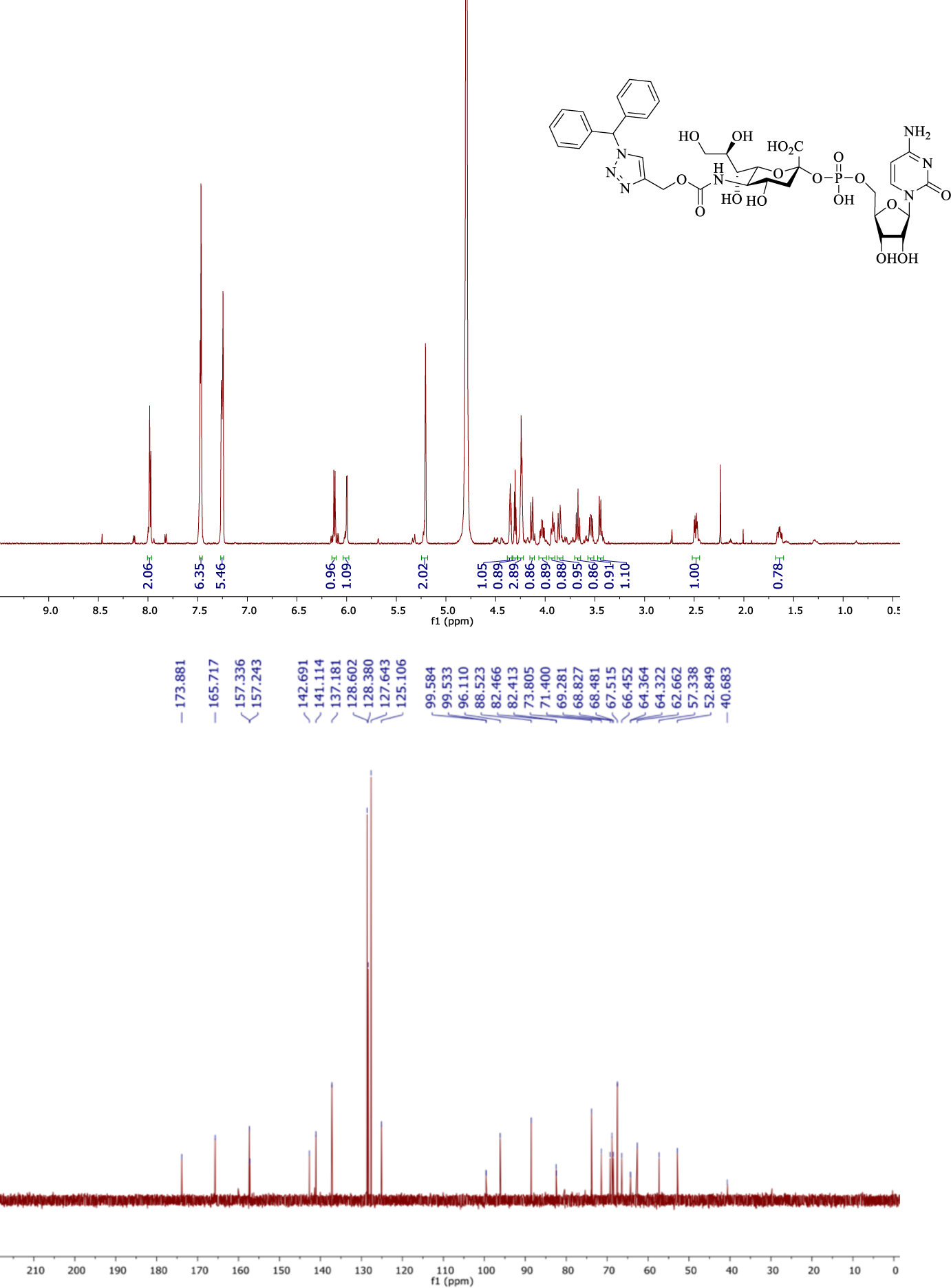
1H and 13C NMR spectra of CMP-^BTMC^Neu5Ac. ^1^HNMR (600 MHz, D_2_O): *δ* = 1.64 (dt, 1H, *J* = 6.0, 12.0 Hz), 2.49 (dd, 1H, *J* = 4.8, 13.2 Hz), 3.45 (d, 1H, *J* = 9.6 Hz), 3.54 (dd, 1H, *J* = 7.2, 12.0 Hz), 3.67 (t, 1H, *J* = 10.8 Hz), 3.86 (dd, 1H, *J* = 2.4, 12.0 Hz), 3.93 (ddd, 1H, *J* = 2.4, 7.2, 9.6 Hz), 4.03 (dt, 1H, *J* = 4.8, 10.8 Hz), 4.14 (d, 1H, *J* = 10.8 Hz), 4.22-4.28 (m, 3H), 4.31 (t, 1H, *J* = 4.8 Hz), 4.35 (t, 1H, *J* = 4.8 Hz), 5.21 (s, 2H), 6.00 (d, 1H, *J* = 4.2 Hz), 6.12 (d, 1H, *J* = 7.8 Hz), 7.22-7.28 (m, 5H), 7.44-7.50 (m, 6H), 7.95-8.01 (m, 2H). ^13^CNMR (150 MHz, D_2_O): *δ* = 40.68, 52.85, 57.34, 62.66, 64.32, 64.36, 66.45, 67.51, 68.48, 68.83, 69.28, 71.4, 73.81, 82.41, 82.47, 88.52, 96.11, 99.53, 99.58, 125.11, 127.64, 128.38, 128.6, 137.18, 141.11, 142.69, 157.24, 157.34, 165.72, 173.88. HRMS: m/z calc. for C_35_H_43_N_7_O_17_P: 864.2453; found 864.2479[M + H]^+^

## Supplementary Information

### Experimental Procedures

#### General methods and materials

All chemical compounds were purchased from commercialized suppliers, and used directly without further purification unless indicated. Az-*PEG*_*4*_-biotin was obtained from Click Chemistry Tools. Streptavidin-Alexa Fluor 647 (AF647) was obtained from Invitrogen. Dynasore, brefeldin A, folic acid, ino-site, 2-me MgSO_4_, ATP, and CTP were purchased from Sigma-Aldrich. Cell culture reagents including media, fetal bovine serum, horse serum, penicillin G, and streptomycin were purchased from GIBCO. The fluorescent images were taken by a Keyence BZ-X800 microscope. The flow cytometry studies were performed on an Attune NxT flow cytometer. Cell Sorting was performed on a Sony multi-application cell sorter MA900.

#### Synthesis of chemical compounds

CMP-Neu5Ac, 9AzSia^[1]^, C9-^CPg^Neu5Ac (**3**)^[2]^, Neu5CPg (**7**)^[3]^, CMP-SiaNAl^[4]^, BTTPS^[5]^ and Ac_4_ManNAl^[6]^ were synthesized and purified following former reports. Enzymes for chemical synthesis, including CMP-Sialic acid synthase (CSS)^[1]^ and inorganic pyrophosphatase (iPPA)^[7]^ were purified from *E. coli* following previously reported procedures.

#### General Procedure for preparation of Sia-CMP

The corresponding unnatural sialic acid (0.1 mmol), along with cytidine triphosphate (0.12 mmol, 1.2 eq) were dissolved in 100 mM Tris and 20 mM MgCl_2_ (3 mL), and the pH was adjusted to 8.5. 10 U of CMP-NeuAc Synthetase was then added to the mixture. The reaction was kept at 37 °C for 2-4 hr until TLC (ethyl acetate: methanol: H_2_O = 5 : 4 : 1 vol/vol) indicated the reaction was complete. The reaction was then quenched with 5 mL methanol for 20 min and centrifuged at 4000 rpm to remove insoluble precipitates. The supernatant was evaporated to remove methanol and the remaining aqueous solution was frozen and lyophilized. After lyophilization, the residue was re-dissolved in H_2_O (1 mL). The solution was then subjected to BTTPS-assisted copper-catalyzed alkyne-azide [3+2] cycloaddition reaction^[5]^ at 30 °C for 4 hr. The solution was then filtered through a 0.22 μm filter and purified through HPLC.

Compound **3** was subjected to the general procedures for the preparation of Sia-CMP and for copper-catalyzed azide-alkyne cycloaddition, which gave compound **5** in a yield of 26% over 2 steps.^1^HNMR (600 MHz, D_2_O): *δ* = 1.84 (t, 1H, *J* = 12.0 Hz), 2.04 (s, 3H), 2.22 (dd, 1H, *J* = 4.8, 12.6 Hz), 3.24 (dd, 1H, *J* = 7.8, 13.8 Hz), 3.47 (d, 1H, *J* = 8.0 Hz), 3.53-3.58 (m, 1H), 3.78-3.84 (m, 1H), 3.89-3.96 (m, 1H), 3.97-4.06 (m, 3H), 4.10-4.15 (m, 1H), 4.24-4.28 (m, 1H), 4.30-4.37 (m, 2H), 5.29-5.37 (m, 2H), 6.00 (d, 1H, *J* = 3.6 Hz), 6.12 (d, 1H, *J* = 7.8 Hz), 6.74-6.83 (m, 4H), 7.25-7.35 (m, 2H), 7.51-7.61 (m, 1H), 7.99-8.08 (m, 2H), 8.27 (s, 1H), 8.65 (s, 1H). ^13^CNMR (150 MHz, D_2_O): *δ* = 22.82, 40.02, 44.42, 52.6, 58.29, 63.91, 67.75, 69.58, 69.86, 69.95, 70.49, 74.68, 83.47, 89.42, 96.66, 103.46, 121.04, 121.7, 122.08, 123.38, 131.61, 131.78, 141.4 (Observed from HSQC). HRMS: m/z calc. for C_44_H_46_N_8_O_22_P: 1069.2464; found: 1069.2487 [M + H]^+^

Compound **7** was subjected to the general procedures for preparation of Sia-CMP and for copper-catalyzed azide-alkyne cycloaddition, which gave compound **9** in a yield of 43% over 2 steps.^1^HNMR (600 MHz, D_2_O): *δ* = 1.64 (dt, 1H, *J* = 6.0, 12.0 Hz), 2.49 (dd, 1H, *J* = 4.8, 13.2 Hz), 3.45 (d, 1H, *J* = 9.6 Hz), 3.54 (dd, 1H, *J* = 7.2, 12.0 Hz), 3.67 (t, 1H, *J* = 10.8 Hz), 3.86 (dd, 1H, *J* = 2.4, 12.0 Hz), 3.93 (ddd, 1H, *J* = 2.4, 7.2, 9.6 Hz), 4.03 (dt, 1H, *J* = 4.8, 10.8 Hz), 4.14 (d, 1H, *J* = 10.8 Hz), 4.22-4.28 (m, 3H), 4.31 (t, 1H, *J* = 4.8 Hz), 4.35 (t, 1H, *J* = 4.8 Hz), 5.21 (s, 2H), 6.00 (d, 1H, *J* = 4.2 Hz), 6.12 (d, 1H, *J* = 7.8 Hz), 7.22-7.28 (m, 5H), 7.44-7.50 (m, 6H), 7.95-8.01 (m, 2H). ^13^CNMR (150 MHz, D_2_O): *δ* = 40.68, 52.85, 57.34, 62.66, 64.32, 64.36, 66.45, 67.51, 68.48, 68.83, 69.28, 71.4, 73.81, 82.41, 82.47, 88.52, 96.11, 99.53, 99.58, 125.11, 127.64, 128.38, 128.6, 137.18, 141.11, 142.69, 157.24, 157.34, 165.72, 173.88. HRMS: m/z calc. for C_35_H_43_N_7_O_17_P: 864.2453; found 864.2479[M + H]^+^

#### Cell culture

Chinese Hamster Ovary (CHO) glycosylation mutant Lec2 cells, NK-92MI cells, Raji cells, Daudi cells, and HEK293T cells were obtained from ATCC and cultured as suggested. Lec2 cells and HEK293T cells were routinely kept in high-glucose DMEM medium supported with 10% (vol/vol) heat-inactivated fetal bovine serum (FBS), 100 U/mL penicillin G, and 100 mg/mL streptomycin. Raji cells and Daudi cells were cultured in RPMI 1640 media supported with 10% (vol/vol) FBS, 10 mM HEPES, 1% (vol/vol) MEM non-essential amino acids, 100 U/mL penicillin G, and 100 mg/mL streptomycin. NK-92MI cells were cultured in αMEM supported with 12.5% FBS, 12.5% horse serum, 200 μM inositol, 20 μM folic acid, 50 μM 2-mercaptoethanol, 100 U/mL penicillin G, and 100 mg/mL streptomycin. The primary NK cells were directly isolated from the blood of healthy donors using a negative isolation kit, following suppler instructions. All individuals provided informed consent for blood donation according to a protocol approved by the Internal Review Board and Ethics Committee. Briefly, a 10 mL blood sample per donor was used to prepare NK cells. The resultant NK cells were collected, washed, and seeded in culture flasks at a concentration of 2 × 10^6^ cells/mL in RPMI-1640 medium (Gibco) containing 10% (vol/vol) FBS (Sigma), MEM NEAA (Gibco), and 10 mM HEPES (Gibco), 100 U/mL penicillin, and 100 U/mL streptomycin (Gibco). The rhIL-2 (100 U/mL) and rhIL-15 (10 ng/mL, Biolegend) were added on day 1. Primary NK cells were kept for 10 days before used for glycocalyx modification and targeting killing assays. Fresh medium, rhIL-2 and rhIL-15 were added every 2 days during culture, and the cell concentration was maintained at 2 × 10^6^ cells/mL. The Siglec-7 high NK-92MI and periphery NK cells were sorted by flow cytometry.

#### Glycan metabolic labelling

The Ac_4_ManNAl was dissolved in ethanol to prepare a 10 mM stock solution. For metabolic labeling of NK-92MI, Raji, or HEK293T cells, Ac_4_ManNAl was preadded to the culture dishes at a final concentration of 50 μM and cells were seeded at a density of 2 × 10^5^ cells/mL after the complete volatilization of ethanol. Cells were then kept in normal culture for 48 h. The resulting SiaNAl-labeled glycoconjugates on cells were further labeled by biotin in PBS buffer (pH 7.4) containing 2% FBS, 50 μM azide-PEG4-biotin, 300 μM BTTPS and 50 μM Cu^2+^ premixture, and 2.5 mM freshly prepared sodium ascorbate for 5 mins. Finally, cell-surface biotin was further probed by 2 μg/mL streptavidin-Alexa fluor 647 conjugate and quantified via flow cytometry. For co-culture treatment, the target cells and effector NK cells were mixed and maintained at 37 °C for 1h before the biotin conjugation.

#### General Procedure for the incorporation of Sia-CMP onto live cells via STs-assisted cell-surface glycan editing

Truncated human sialyltransferases were prepared as previously described^[8]^, including hST6Gal1, hST3Gal1, and hST3Gal4. For STs-assisted cell-surface glycan editing, the live cells were washed three times with PBS buffer (pH 7.4) and resuspended in HBSS buffer (pH 7.4) containing 3 mM HEPES, 20 mM MgSO_4_, 40 μg/mL hSTs, and indicated concentrations of CMP-Sia. The Siglec staining buffer was prepared by premixing the Siglec-7 Fc or Siglec-9 Fc (5 μg/mL) with anti-human Fc APC conjugate at a 1:1.2 molar ratio on ice for 1 hr. hSTs-edited cells were washed with PBS containing 1% (vol/vol) FBS for three times, resuspended in Siglec staining buffer and incubated for 30 min on ice in the dark. Then, the cells were washed and resuspended in FACS buffer for flow cytometry analysis. For fluorescence microscopy imaging, hSTs-edited NK-92MI cells were washed with PBS containing 1% (vol/vol) FBS for three times and incubated with Raji target cells at an E/T ratio of 5:1 on glass-bottom dishes (35 mm) for 1 hr at 30 °C. The resulting cell-cell clusters were stained in 1:1 immunoblot buffer (PBS, pH 7.4, 2 % (vol/vol) FBS, 2 μg/mL antiSiglec-7 PE conjugate) Siglec staining buffer, and incubated for 30 min on ice in dark. Finally, the cells were carefully washed with PBS twice and imaged via fluorescence microscopy.

#### LDH assay

NK-92MI cell cytotoxic function was evaluated by LDH assays using Raji and Daudi B lymphoma cells as targets for cytotoxicity. The number of effector cells available was determined via hemocytometer, and the viability of cells was assessed by trypan blue exclusion. Effector cells and target cells were used as controls for assays. The counts of periphery NK cells were determined via flow cytometry by gating anti-CD3-FITC (negative) and anti-CD56-PE (positive). The LDH assays were performed by following the instructions from the commercial supplier. In brief, E/T coculture of the indicated cells in about 100 μL media was maintained at 37 °C for 4∼5 hr, and 40 μL supernatant was collected and subjected to LDH assay. The pretreatment of NK cells with Siglec-7 blocking antibody (clone s7.7, 2.5 μg/mL, Biolegend) was performed by incubation cells at 37 °C for 30 min, and α2-3,6,8,9-neuraminidase A (NEB) assisted desialylation of NK cells or target cells was performed by incubating cells at 37 °C for 60 min, before coincubation and target cell killing assays.

#### Flow cytometry assays

The Siglec-7 on NK-92MI cells were stained with mouse anti-Siglec-APC antibody on ice for 30 min in dark (clone 6-434, 1:200, Biolegend). The Siglec-7 ligands were detected with the premixture (1h on ice in dark) of Siglec-7 Fc and anti-human Fc-APC antibody on ice for 30 min in dark (1:200, Biolegend). Lectin stainings were performed with SNA-biotin (10 μg/mL, Vector) and MAA-II-biotin (10 μg/mL, Vector) on ice for 30 min in dark, following streptavidin-Alexa Fluor 647 (2 μg/mL, Life) after three times washing with DPBS (pH 7.4). In the coculture system, the NK-92MI cells were detected via anti-CD56-PE (1:200, Biolegend), periphery NK cells were probed by anti-CD3-FITC (negative) and anti-CD56-PE (positive), the Raji cells were detected with anti-CD19-APC (1:200, Biolegend). After staining, the cells were spun down, washed with DPBS for three times, and resuspended in FACS buffer for flow cytometry assays.

#### Fluorescence microscopy imaging

For imaging of live cells, the NK-92MI cells modified with or without Siglec-7 specific high-affinity ligands were further incubated with Raji target cells at a 5:1 E/T ratio for 1h at 37 oC, or not. Then, the Siglec-7 on NK-92MI cells were stained directly by mouse anti-Siglec-7-PE antibody (clone 6-434, 1:200, Biolegend). The signal of PE and fluorescein was directly imaged after washing off the free dyes. The neuraminidase assisted desialylation of NK cells was performed by incubation cells at 37 °C for 60 min, before coincubation with or without target Raji cells. For immunostaining of the tissue specimens, the paraffin-embedded human tumor and adjacent tissue specimens from a number of malignancies were purchased from commercial suppler (US Biomax), including MC245a (c040 and c041), lung tissue array BC041114 (E094 and E095) and (LC1201a). The Histo-clear II (Electron Microscopy Sciences) was used to remove the paraffin following the supplier instructions. In brief, the slides were washed in Histo-clear II for 2 times (5 min), 10% ethanol (vol:vol) in Histo clear II for 1 time (5 min), ethanol for 2 times (5 min), 95% ethanol (vol:vol) in H_2_O for 1 time (5 min), 80% ethanol (vol:vol) in H_2_O for 1 time (5 min), and 70% ethanol (vol:vol) in H_2_O for 1 time (5 min). The tissue samples were then blocked by incubation with DPBS buffer (pH7.4) containing 5% (w/vol) bovine serum albumin (BSA) at rt for 1h. After washed with DPBS three times, the slides were further stained with antibodies by incubation at 4°C overnight. To probe the sialyl-ligands of Siglec-7 on tissuses, the Siglec-7 Fc (2 μg/mL) were premixed with anti-human IgG Fc-APC conjugated monoclonal antibody (5 μg/mL, Biolegend) at 4 °C for 30 min, before added to tissue samples. The NK cells in tissue samples were probed by the human CD56 antibody (2 μg/mL, Biolegend). Then, the slides were washed with DPBS 3 times, mounted with the prolonged gold antifade media with DAPI (Cell Signaling), and sealed for fluorescent imaging.

#### Western blot and immunoprecipitation (IP)

The NK-92MI-S7^high^ cells (∼10 × 10^6^) modified with or without ^FTMC^Neu5Ac were mixed with target cells in RPMI 1640 medium (100 uL) at the indicated effector to target ratio for 30 min or 1h and pelleted down. The cells were washed with DPBS and lysed with RIPA buffer containing 5 μg/mL DNase 1 and protease inhibitor cocktail. For Siglec-7 IP, the resultant lysates were first incubated with 10 μg Rabbit anti-Siglec-7 antibody (clone H-48, Santa Cruz Biotech) for 3h, followed by protein A/G agarose incubation overnight at 4 °C. The beads were washed three times with the lysis buffer and eluted by boiling in 4 x SDS loading buffer containing β-mercaptoethanol. For Western blotting, proteins were resolved by SDS-PAGE on Bis-Tris Criterion Gels (4-12%; Bio-Rad) and transferred to nitrocellulose membrane by wet transfer (Tris-glycine, 20% MeOH) at 100 V for 1h. the membranes were blocked with PBST buffer (DPBS with 0.05% Tween-20) containing 5% BSA, and the primary antibody incubation conditions were conducted in PBST buffer containing 1% BSA overnight. The antibodies used in this study were mouse anti-Siglec-7 antibody (clone 194212, 1:500, R&D)/HRP goat anti-mouse IgG (1:10,000, Pierce®), biotin anti-phosphotyrosine (clone PY20, 1:200, Biolegend)/HRP anti-biotin antibody (1:10,000, Jackson), rat anti-SHP-1 antibody (clone W17240D, 1:500, Biolegend)/HRP goat anti-rat IgG antibody (1:5000, Biolegend).

#### Data and software availability

The fluorescent images were processed with ImageJ and the flow cytometry data was processed using Flowjo. The raw data that supported the findings of this study are available from the authors upon reasonable request.

## References

(1) Miller, J. F.; Sadelain, M. The journey from discoveries in fundamental immunology to cancer immunotherapy. Cancer Cell 2015, 27, 439–449.

(2) Ribas, A.; Wolchok, J. D. Cancer immunotherapy using checkpoint blockade. Science 2018, 359, 1350–1355.

(3) Sharma, P.; Allison, J. P. The future of immune checkpoint therapy. Science. 2015, 348, 56–61.

(4) Barkal, A. A. et al. CD24 signalling through macrophage Siglec-10 is a target for cancer immunotherapy, Nature 2019, 572, 392–396.

(5) Wang, J. et al. Siglec-15 as an immune suppressor and potential target for normalization cancer immunotherapy, Nat. Med. 2019, 25, 656–666.

(6) Jandus, C. et al. Interactions between Siglec-7/9 receptors and ligands influence NK cell–dependent tumor immunosurveillance, J. Clin. Invest. 2014, 124, 1810–1820.

(7) Hudak, J. E.; Canham, S. M.; Bertozzi, C. R. Glycocalyx engineering reveals a Siglec-based mechanism for NK cell immunoevasion, Nat. Chem. Biol. 2014, 10, 69–75.

(8) Macauley, M. S.; Crocker, P. R.; Paulson, J. C. Siglec-mediated regulation of immune cell function in disease, Nat. Rev. Immunol. 2014, 14, 653–666

(9) Rodriguez, E.; Schetters, S. T. T.; Kooyk, Y. V. The tumour glyco-code as a novel immune checkpoint for immunotherapy, Nat. Rev. Immunol. 2018, 18, 204–211

(10) Ikehara, Y.; Ikehara, S. K.; Paulson, J. C. Negative regulation of T cell receptor signaling by Siglec-7 (p70/AIRM) and Siglec-9, J. Biol. Chem. 2004, 279, 43117–43125.

(11) Kawasaki, Y. et al. Ganglioside DSGb5, preferred ligand for Siglec-7, inhibits NK cell cytotoxicity against renal cell carcinoma cells. Glycobiology 2010, 20, 1373–1379.

(12) Tao, L.; Wang, S.; Yang, L.; Jiang, L.; Li, J.; Wang, X. Reduced Siglec-7 expression on NK cells predicts NK cell dysfunction in primary hepatocellular carcinoma, Clin. Exp. Immunol. 2020, DOI: 10.1111/cei.13444

(13) Laubli, H. et al. Engagement of myelomonocytic Siglecs by tumor-associated ligands modulates the innate immune response to cancer. Proc. Natl. Acad. Sci. USA 2014, 111, 14211–14216.

(14) Adams, O. J.; Stanczak, M. A.; von Gunten, S.; Läubli, H.; Targeting sialic acid–Siglec interactions to reverse immune suppression in cancer. Glycobiology 2018, 28, 640–647.

(15) Bornhofft, K. F.; Goldammer, T.; Rebl, A.; Galuska, S. P.; Siglecs: A journey through the evolution of sialic acid-binding immunoglobulin-type lectins. Dev. Comp. Immunol. 2018, 86, 219–231.

(16) Vivier, E. et al. Innate or adaptive immunity? The example of natural killer cells. Science 2011, 331, 44–49.

(17) Veluchamy, J. P. et al. The Rise of Allogeneic Natural Killer Cells As a Platform for Cancer Immunotherapy: Recent Innovations and Future Developments. Frontiers in immunology 2017, 8, 631.

(18) Suck, G. et al. NK-92: an ‘off-the-shelf therapeutic’ for adoptive natural killer cell-based cancer immunotherapy. Cancer Immunol. Immunother. 2016, 65, 485–492.

(19) Zhu, L. et al. Natural Killer Cell (NK-92MI)-Based Therapy for Pulmonary Metastasis of Anaplastic Thyroid Cancer in a Nude Mouse Model. Frontiers in immunology 2017, 8, 816.

(20) Hydes, T. et al. IL-12 and IL-15 induce the expression of CXCR6 and CD49a on peripheral natural killer cells. Immun. Inflamm. Dis. 2018, 6, 34–46.

(21) Carson, W. E. et al., Interleukin (IL) 15 is a novel cytokine that activates human natural killer cells via components of the IL-2 receptor. J. Exp. Med. 1994, 180, 1395–1403.

(22) Gong, J. H.; Maki, G.; Hlingemann, H. G. Characterization of a Human Cell Line (NK-92) With Phenotypical and Functional Characteristics of Activated Natural Killer Cells. Leukemia 1994, 8, 652–658.

(23) Wratil, P. R. et al. Metabolic glycoengineering with N-acyl side chain modified mannosamines, Angew Chem. Int. Ed. Engl. 2016, 55, 9482–9512.

(24) Wang, W. et al. Sulfated ligands for the copper(I)-catalyzed azide-alkyne cycloaddition. Chem. Asian J. 2011, 6, 2796–2802.

(25) Donaldson, J. G.; Finazzi, D.; Klausner, R. D. Brefeldin A inhibits Golgi membrane-catalysed exchange of guanine nucleotide onto ARF protein, Nature 1992, 360, 350–352.

(26) Macia, E.; Ehrlich, M.; Massol, R.; Boucrot, E.; Brunner, C.; Kirchhausen, T. Dynasore, a cell-permeable inhibitor of dynamin. Dev. Cell 2006, 10, 839–850.

(27) Zhou, Z. et al. Granzyme A from cytotoxic lymphocytes cleaves GSDMB to trigger pyroptosis in target cells, Science 2020, 368, eaaz7548.

(28) Avril, T.; North, S. J.; Haslam, S. M.; Willison, H. J.; Crocker, P. R. Probing the cis interactions of the inhibitory receptor Siglec-7 with alpha2,8-disialylated ligands on natural killer cells and other leukocytes using glycan-specific antibodies and by analysis of alpha2,8-sialyltransferase gene expression. J. Leukoc. Biol. 2006, 80, 787–796.

(29) Nicoll, G.; Avril, T.; Lock, K.; Furukawa, K.; Bovin, N.; Crocker, P. R. Ganglioside GD3 expression on target cells can modulate NK cell cytotoxicity via siglec-7-dependent and -independent mechanisms. Eur. J. Immunol. 2003, 33, 1642–1648.

(30) Rillahan, C. D. et al. On-chip synthesis and screening of a sialoside library yields a high affinity ligand for Siglec-7. ACS Chem. Biol. 2013, 8, 1417–1422.

(31) Sugiarto, G. et al. A sialyltransferase mutant with decreased donor hydrolysis and reduced sialidase activities for directly sialylating LewisX. ACS Chem. Biol.. 2012, 7, 1232–1240.

(32) Briard, J. G.; Jiang, H.; Moremen, K. W.; Macauley, M. S.; Wu, P. Cell-based glycan arrays for probing glycan-glycan binding protein interactions. Nat. Commun. 2018, 9, 880–890.

(33) Patnaik, S. K.; Stanley, P. Lectin-resistant CHO glycosylation mutants. Methods Enzymol. 2006, 416, 159–182.

(34) Aguilar, A. L.; Briard, J. G.; Yang, L.; Ovryn, B.; Macauley, M. S.; Wu, P. Tools for studying glycans: recent advances in chemoenzymatic glycan labeling. ACS Chem. Biol. 2017, 12, 611–621.

(35) Moremen, K. W. et al. Expression system for structural and functional studies of human glycosylation enzymes. Nat. Chem. Biol. 2018, 14, 156–162.

(36) Collins, B. E.; Smith, B. A.; Bengtson, P.; Paulson, J. C. Ablation of CD22 in ligand-deficient mice restores B cell receptor signaling. Nat. Immunol. 2006, 7, 199–206.

(37) Wherry, E. J.; Kurachi, M. Molecular and cellular insights into T cell exhaustion. Nat. Rev. Immunol. 2015, 15, 486–499.

(38) Blank, C. U. et al. Defining ‘T cell exhaustion’. Nat. Rev.Immunol. 2019, 19, 665–674.

(39) Spranger, S. et al. Up-regulation of PD-L1, IDO, and T_regs_in the melanoma tumor microenvironment is driven by CD8^+^ T cells. Sci. Trans. Med. 2013, 5, 200ra116.

(40) Varchetta, S. et al. Lack of Siglec-7 expression identifies a dysfunctional natural killer cell subset associated with liver inflammation and fibrosis in chronic HCV infection. Gut, 2015, 0, 1–9.

(41) Rillahan, C. D.; Schwartz, E.; McBride, R.; Fokin, V. V.; Paulson, J. C. Click and pick: identification of sialoside analogues for siglec-based cell targeting. Angew Chem. Int. Ed. Engl. 2012, 51, 11014–11018.

## References

[1] H. Yu, H. Yu, R. Karpel, X. Chen, Chemoenzymatic synthesis of CMP-sialic acid derivatives by a one-pot two-enzyme system: comparison of substrate flexibility of three microbial CMP-sialic acid synthetases, Bioorg. Med. Chem. 2004, 12, 6427–6435.

[2] C. D. Rillahan, E. Schwartz, R. McBride, V. V. Fokin, J. C. Paulson, Click and pick: identification of sialoside analogues for siglec-based cell targeting, Angew. Chem. Int. Ed. Engl. 2012, 51, 11014–11018.

[3] Y. Li, H. Yu, H. Cao, S. Muthana, X. Chen, Pasteurella multocida CMP-sialic acid synthetase and mutants of Neisseria meningitidis CMP-sialic acid synthetase with improved substrate promiscuity, Appl. Microbiol. Biotechnol. 2012, 93, 2411–2423.

[4] M. Noel, P. A. Gilormini, V. Cogez, N. Yamakawa, D. Vicogne, C. Lion, C. Biot, Y. Guérardel, A. Harduin-Lepers, Probing the CMP-sialic acid donor specificity of two human β-D-galactoside sialyltransferases (ST3GalI and ST6GalI) selectively acting on O-and N-glycosylproteins Chembiochem 2017, 18, 1251–1259.

[5] W. Wang, S. Hong, A. Tran, H. Jiang, R. Triano, Y. Liu, X. Chen, P. Wu, Sulfated ligands for the copper(I)-catalyzed azide-alkyne cycloaddition, Chem Asian J 2011, 6, 2796–2802.

[6] P. V. Chang, X. Chen, C. Smyrniotis, A. Xenakis, T. Hu, C. R. Bertozzi, P. Wu, Metabolic labeling of sialic acids in living animals with alkynyl sugars, Angew. Chem. Int. Ed. Engl. 2009, 48, 4030–4033.

[7] W. Wang, T. Hu, P. A. Frantom, T. Zheng, B. Gerwe, D. S. del Amo, S. Garret, R. D. Seidel, P. Wu, Chemoenzymatic synthesis of GDP-L-fucose and the Lewis X glycan derivatives, Proc. Natl. Acad. Sci. U.S.A. 2009, 106, 16096–16101.

[8] K. W. Moremen, A. Ramiah, M. Stuart, J. Steel, L. Meng, F. Forouhar, H. A. Moniz, G. Gahlay, Z. Gao, D. Chapla, et al., Human glycosylation enzymes for enzymatic, structural and functional studies, Nat. Chem. Biol. 2018, 14, 156–162.

